# Differential detection workflows for multi-sample single-cell RNA-seq data

**DOI:** 10.1101/2023.12.17.572043

**Authors:** Jeroen Gilis, Laura Perin, Milan Malfait, Koen Van den Berge, Alemu Takele Assefa, Bie Verbist, Davide Risso, Lieven Clement

## Abstract

In single-cell transcriptomics, differential gene expression (DE) analyses typically focus on testing differences in the average expression of genes between cell types or conditions of interest. Single-cell transcriptomics, however, also has the promise to prioritise genes for which the expression differ in other aspects of the distribution. Here we develop a workflow for assessing differential detection (DD), which tests for differences in the average fraction of samples or cells in which a gene is detected. After benchmarking eight different DD data analysis strategies, we provide a unified workflow for jointly assessing DE and DD. Using simulations and two case studies, we show that DE and DD analysis provide complementary information, both in terms of the individual genes they report and in the functional interpretation of those genes.

## 1. Introduction

Single-cell RNA-sequencing (scRNA-seq) has improved our understanding of complex biological processes by elucidating cell-level heterogeneity in gene expression. One of the key tasks in the downstream analysis of scRNA-seq data is studying differential gene expression (DE). Traditional DE analyses aim to identify genes for which the average expression differs between biological groups of interest, e.g., between cell types or between diseased and healthy cells. Unfortunately, traditional DE analyses only allow for assessing one aspect of the gene expression distribution: the mean. However, in scRNA-seq data, other differences between count distributions can be observed, e.g. differences in the number of modes and differential variability. This has recently prompted the development of a variety of frameworks that allow for comparing multiple aspects of the expression distribution (Korthauer et al., 2016, Zhang et al. (2022), Tiberi et al. (2023)).

One particularly interesting characteristic of gene expression not explicitly captured by the aforementioned frameworks is differential detection (DD), i.e. finding differences in the fraction of cells in which a gene is detected between groups. It has been reported that gene expression profiles may exhibit characteristic bimodal expression patterns, in which the expression of genes is either strongly positive or undetected within individual cells (e.g., Finak et al., 2015). While differences in gene detection may arise from technical artifacts or from the stochastic nature of gene expression (Hicks et al., 2018), they can also reflect biologically meaningful differences between samples. In this respect, Qiu (2020) has demonstrated that binarizing scRNA-seq counts generates expression profiles that still accurately reflect biological variation. This was confirmed by Bouland et al. (2021), who showed that the frequencies of zero counts capture biological variability and even claimed that a binarized representation of the single-cell expression data allows for a more robust description of the relative abundance of transcripts than counts.

MAST (Finak et al., 2015) was one of the first methods for jointly studying differences in gene detection and gene expression. It adopts a two-component generalized linear model, commonly referred to as a Hurdle model. The first component consists of a logistic regression model on the binarized expression matrix, which can make inference on differential detection between conditions. The second component models the gene expression of cells for which the gene has positive counts, using a Gaussian model on log-transformed and normalized counts, and can be used to infer differential expression (DE) given that the gene is detected in a cell. Furthermore, Finak et al. (2015) use the cellular detection rate (CDR), i.e., the fraction of genes detected in each cell, as an additional covariate in both model components to normalize for between-cell differences due to technical variability, e.g. mRNA quality, pre-amplification efficiency of the scRNA-seq assay, and extrinsic biological factors, e.g. nuisance biological variability such as differences in cell volume (Finak et al., 2015). For the inference, MAST can adopt either a Wald test or a likelihood ratio test, building on the conditional independence of both parts of the hurdle model.

MAST models the data directly at the single-cell level. This poses several challenges. First, its scalability is compromised due to the large increase in the number of cells that are being profiled in modern scRNA-seq experiments. Second, as discussed in Townes et al. (2019), the sparsity of the data of popular droplet-based protocols renders the inference with Gaussian models on log-transformed counts invalid. The extensive scRNA-seq differential expression benchmark of Soneson and Robinson (2018) confirmed that MAST did not outperform classical bulk RNA-seq analysis tools such as edgeR (Robinson et al., 2010), DESeq2 (Love et al., 2014) and limma (Smyth, 2004). Thirdly, several studies have highlighted that gene expression in multi-sample/multi-cell experiments exhibit a complex correlation pattern. Indeed, the expression of cells from the same sample is more similar than that of cells across samples and this within-sample correlation should be addressed in the statistical inference (Lun and Marioni, 2017, Zimmerman et al. (2021) and Squair et al. (2021)).

Zimmerman et al. (2021) and Murphy and Skene (2022) showed that this correlation structure could be addressed in MAST by including a sample-specific random intercept. However, this further increases MAST’s computational burden. An alternative strategy to address the within sample correlation is to perform pseudobulk aggregation (e.g., (Lun and Marioni, 2017, and Crowell et al. (2020))). Crowell et al. (2020) proposed summing the gene expression counts for cells within the same cell type-sample combination to obtain pseudobulk samples, and subsequently modeling the aggregated counts via negative binomial generalized linear models (GLMs) using edgeR. In their benchmarks, this strategy outperformed MAST without random effects and performed similarly to mixed-effect GLMs. Moreover, Murphy and Skene (2022) evaluated several proposed workflows based on MAST, including MAST with random effects, and showed that all of these were outperformed by running edgeR on pseudobulk aggregated data.

In this work, we show the potential of differential detection (DD) strategies for scRNA-seq data analysis of multi-sample/multi-cell experiments. First, we benchmark eight different DD strategies: we start with a simple binomial regression model on the binarized scRNA-seq expression matrix, and gradually increase the model complexity to account for overdispersion and allow for model-based normalization using the CDR as proposed by (Finak et al., 2015). Pseudobulking the binarized single cell counts is a natural strategy in the context of multi-sample/multi-cell datasets; it improves model performance, type I error control and tremendously decreases the computational complexity compared to a single-cell level analysis. Moreover, our workflow allows for assessing differential detection and differential expression simultaneously. Inference on both hypotheses can be combined using the two-stage testing paradigm of Van den Berge et al. (2017). Finally, we show the added value of our two-stage test for DD and DE on two large multi-sample case studies.

## 2. Results

Multi-sample/multi-cell scRNA-seq experiments imply a hierarchical correlation structure of the data at the single cell level. The state-of-the-art methods for assessing DE that properly address this correlation structure rely on pseudobulk strategies that first aggregate single cell counts by taking their sum at the sample-level for each cell type, which can then be analyzed using conventional bulk RNA-seq methods (e.g. Crowell et al. (2020)). The frequencies of zero counts, however, also capture biological variability (e.g. (Finak et al., 2015, and Bouland et al. (2021))) and a binarized representation of the single-cell RNA-seq counts allows to infer DD. Pseudobulk strategies for inferring DD are also very natural. Indeed, upon summing the binary counts of cells from the same sample for each cell type, a binomial distribution with the total number of cells for a sample as the “number of trials” and the proportion of cells that express the gene as “success probability” is obtained. Pseudobulking of binarized counts also effectively addresses the within-sample correlation and dramatically reduces the computational complexity. Here, we develop eight DD workflows, four workflows based on binomial regression models and four workflows that model the aggregated binomial counts using edgeR. We refer the reader to Section 4 for more details.

The results section is organized as follows: we first benchmark the type I error control of the eight DD workflows in null simulations, their sensitivity and specificity in simulations with know ground truth, and the effect of sample size on their performance. Next, we show how combining results from DD and DE analyses can increase the power to detect differential genes, and allows for pinpointing if the changes in expression are due to differences in detection, differences in expression, or both. Finally, we use the best performing DD workflow and combine it with a conventional pseudobulk DE analysis in two case studies on publicly available datasets from large multi-sample studies.

### 2.1. Mock comparison

We benchmark the type I error control of the eight proposed DD workflows in mock simulations where none of the genes are differential. All genes that are flagged as DD between conditions are thus false positives. The mock simulation datasets were generated based on two real single-cell transcriptomics datasets, here referred to as the “lupus” and “COVID” datasets, obtained from Perez et al. (2022) and Stephenson et al. (2021), respectively. We refer the reader to Section 4.4 for more details on the design of the mock simulation studies.

In Figure 1, we show the p-value densities of the eight DD workflows for five replicate mock simulations. The mock simulation is based on the non-classical myeloid cells from the lupus dataset and consists of five samples in each mock treatment group after binarization and pseudobulk aggregation. The analysis was performed five times, each time with a different randomly selected subset of five samples per mock treatment group. For these mock simulations, the p-value densities will follow the uniform distribution when the DD method is controlling the type I error as desired, which would appear in Figure 1 as horizontal lines. The inference of the regular binomial GLM (bGLM) is overly liberal across all five simulation replicates, which can be seen by the large peaks for small p-values in Figure 1, panel A. By allowing for over- and underdispersion with respect to the binomial variance, the quasi-binomial GLM (qbGLM) produces more uniformly distributed p-values (Figure 1, panel B). The type I error control can be further improved by including a normalization offset (Figure 1, panels C and D). The two baseline edgeR-based workflows, edgeR_NB and edgeR_QP are generally conservative and display a peak of p-values equal to 1 (Figure 1, panels E and F). By removing genes that are detected in at least 90% of cells in each pseudobulk sample, incorporating a normalization offset and allowing for underdispersion, the edgeR_NB_optim and edgeR_QP_optim workflows also provide a proper type I error control (Figure 1, panels G and H).

**Figure 1:**
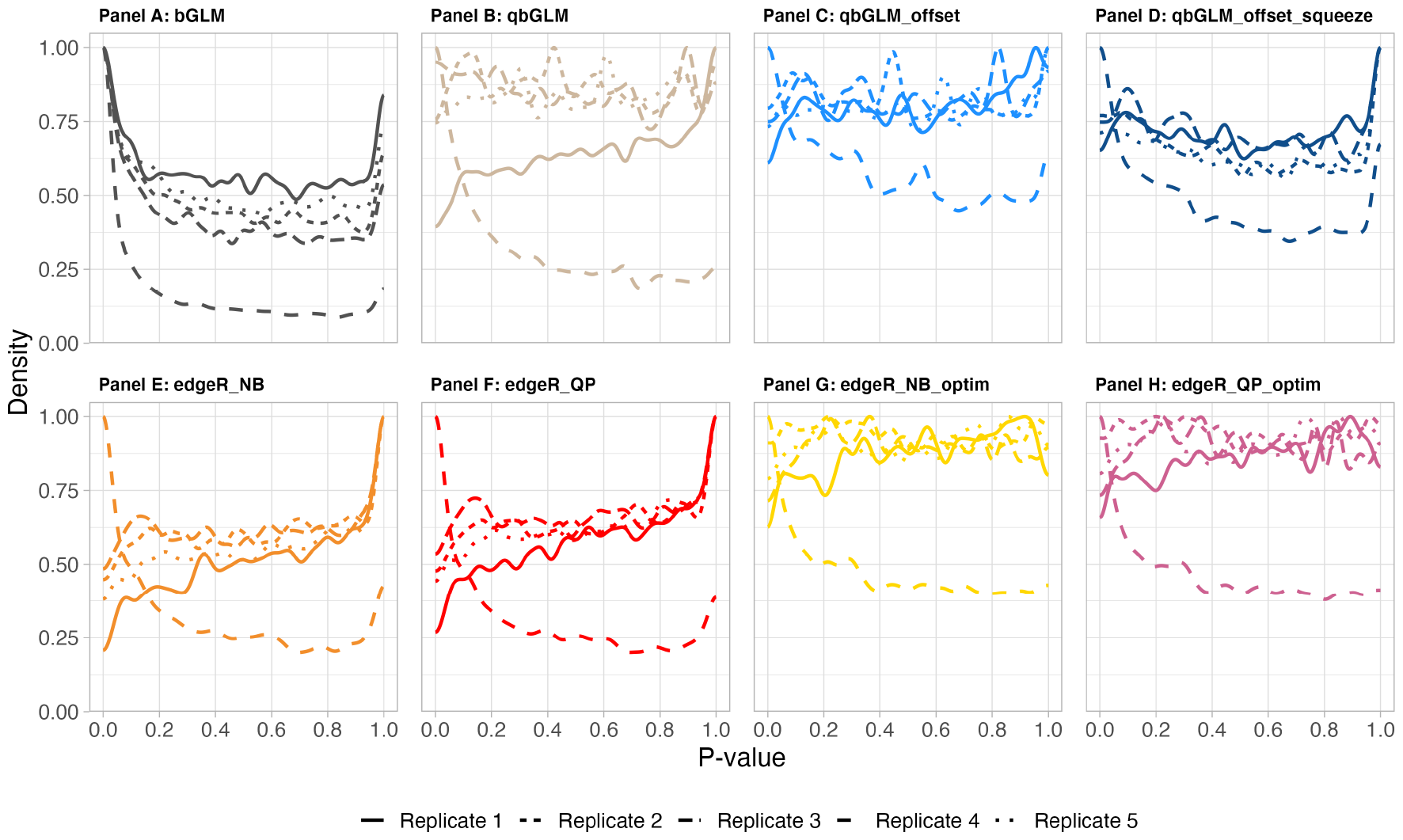
Nominal p-value densities obtained from five null simulation replicates, stratified by method. The null simulations are based on the non-classical myeloid cells from the lupus dataset and are binarized and aggregated to pseudobulk samples, thus creating a 5 versus 5 comparison. The different line types indicate the p-value densities for the different replicates. **If a DD method accurately controls the type I error, the p-value densities will follow the uniform distribution, which will appear in the visualization as horizontal lines**.

In *Supplementary Figures S1-S3*, we show the p-value densities of the eight methods on twelve null simulated datasets with five replicates each, consisting of four different sample sizes (5 verus 5, 10 versus 10, 15 versus 15 and 22 versus 22 samples after pseudobulk aggregation) and based on three different cell types of the lupus dataset, i.e., T4 naive cells, B memory cells and non-classical myeloid cells (see Section 4.4). The p-value densities obtained for these twelve datasets are in line with those displayed in Figure 1. Note, that we do not observe pronounced effects of sample size on the type I error control of the different methods.

When analyzing the same datasets of 22 versus 22 patients on the single-cell level, i.e., without pseudobulk aggregation, the results of all methods become overly liberal (*Supplementary Figure S4*). This is in line with previous observations that cells from the same sample are correlated, and that the within-sample correlation should be addressed accordingly in the statistical inference (Lun and Marioni, 2017, Zimmerman et al. (2021) and Squair et al. (2021)). We also note that the inference is most liberal for the T4 naive cell type, which has the most cells per patient in this dataset.

In the null simulations based on the COVID dataset, differences between the eight methods are more pronounced (*Supplementary Figures S5-S7*). All binomial GLM-based methods display a peak of p-values equal to 1. These correspond to genes for which the overall detection, i.e., the fraction of cells in each pseudobulk sample that express the gene, is low. While the same gene-level filtering strategy was adopted when generating the lupus and COVID simulation datasets (see Section 4), such sparse genes are more common in the COVID dataset. The regular binomial GLM additionally has a large peak of p-values near zero, which is consistent with the observations in Figure 1 panel A that the binomial GLM does not control the type I error at the nominal level. Using quasi-binomial models and additionally including a normalization offset improves the type I error control, however, the peaks of p-values equal to 1 are still present. Across the different cell types, edgeR-based methods generally display a reasonable control of the type I error, with the egdeR_optim workflows performing better than the default edgeR methods. Note, that none of the edgeR-based methods display peaks of p-values equal to 1. This is likely because edgeR internally adds a small count to the data, called a pseudo-count, which is used for stabilizing the parameter estimation procedure when data is sparse and unties the zero counts when the pseudobulk samples differ in their number of cells.

### 2.2. Performance evaluation

We assessed the performance of the eight DD workflows on simulated datasets with known ground truth. As in the previous paragraph, the simulated datasets were generated based on the lupus dataset by Perez et al. (2022) and the COVID dataset by Stephenson et al. (2021); however, in this case the simulation framework allows to have DD genes and we evaluate the workflows both in terms of type I error control and power. We refer to Section 4.6 for more details on the setup of these simulation studies.

In Figure 2, we show the benchmark results on simulated data based on three cell types from the lupus dataset, i.e., non classical myeloid (ncM) cells, T4 naive cells and memory B cells. The simulated data for each cell type consists of 5 samples in each mock treatment group after binarization and pseudobulk aggregation. The analysis was performed five times per cell type, each time with a different randomly selected subset of five samples per treatment group, and results were averaged across replicate analyses. Each curve in Figure 2 visualizes the performance of a DD workflow by displaying its sensitivity (true positive rate, TPR) versus its specificity (false discovery rate, FDR). We refer to Section 4.7 for a detailed description of how these measure were computed. All edgeR-based methods consistently outperform the binomial GLM-based methods. For the simulations based on the ncM cell type, the cell type with the smallest average number of cells per patient, the edgeR_optim strategies outperform the baseline edgeR workflows. For all datasets, the inference of the canonical binomial regression model is too liberal. Adopting a quasi-binomial model improves the type I error control. Incorporating a normalization offset in the quasi-binomial model and additionally shrinking the dispersion estimates increases the power of the binomial GLM-based workflows.

**Figure 2:**
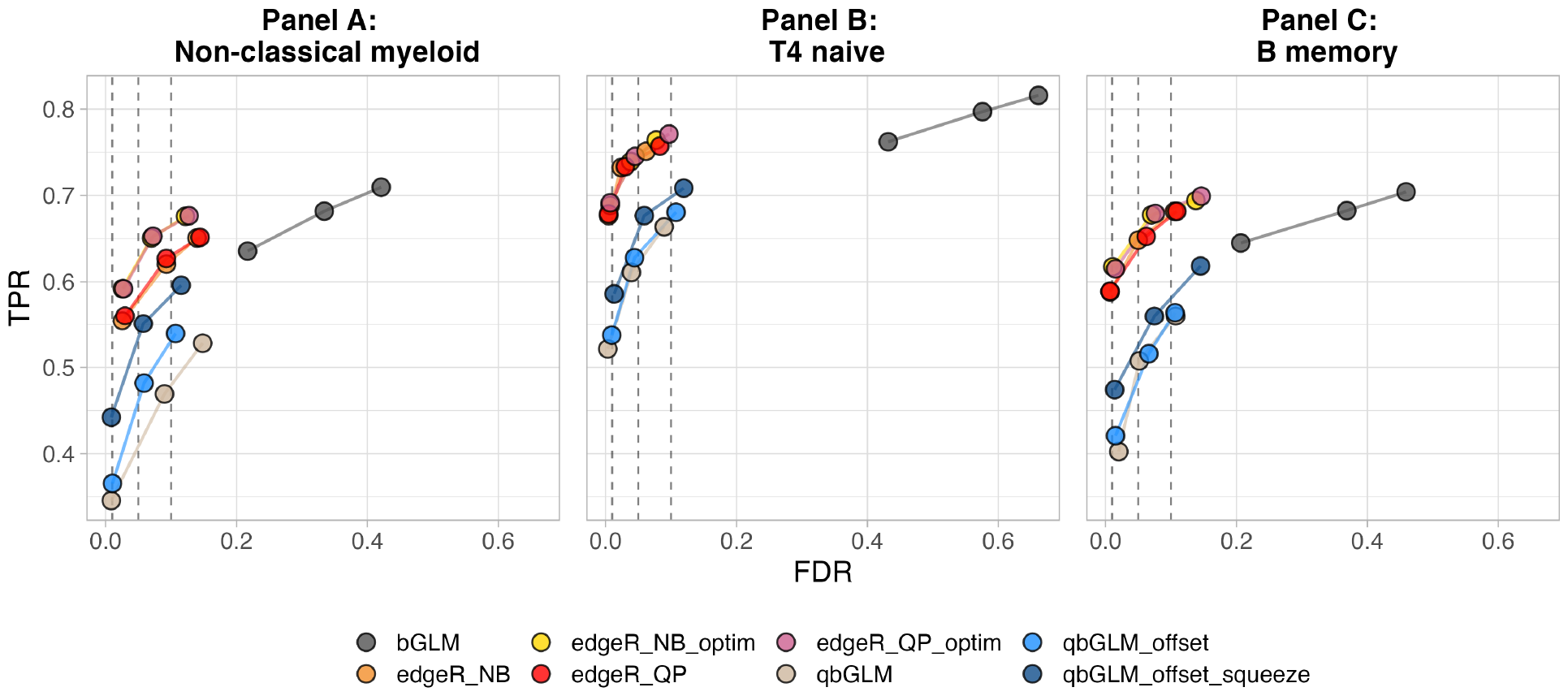
Performance evaluation of eight differential detection workflows on three simulated datasets. The simulated data is based on the three different celltypes (non-classical myeloid cells, T4 naive cells and memory B cells) of the lupus dataset. The data was binarized and aggregated to the pseudobulk level, resulting in a test between two groups of five samples each. Each curve visualizes the performance of each method by displaying the sensitivity of the method (true positive rate, TPR) with respect to the false discovery rate (FDR). The curves display averages over 5 replicates for each simulated dataset. The three circles on each curve represent working points when the FDR level is set at nominal levels of 1%, 5% and 10%, respectively.

*Supplementary Figure S8* provides an alternative visualization for the performance benchmark on the same data used to construct Figure 2. Here, the false discovery proportion (FDP) and true positive proportion (TPP) of each method are displayed for each individual dataset replicate. Except for the canonical binomial GLM being overly liberal, the observed FDP of the other methods is close to the nominal level for all replicates.

*In Supplementary Figure S9*, we show the performance benchmark results on the same three cell types from the lupus dataset for four different sample sizes (5 versus 5, 10 versus 10, 15 versus 15 and 22 versus 22 samples after pseudobulk aggregation). As expected, the power increases with increasing sample size. edgeR-based methods outperform binomial GLM-based methods across all sample sizes. Note, that the canonical binomial GLM was excluded from these plots, because the inference remained overly liberal across all benchmarks, obscuring the differences between the other methods.

*Supplementary Figure S10* shows the results from the performance benchmark on simulated data generated based on the COVID dataset. Again, all edgeR-based methods consistently outperform the binomial GLM-based methods. For all datasets, the inference of the canonical binomial regression model is overly liberal. The quasi-binomial model and the quasi-binomial model with a normalization offset have very poor performances on some of the datasets, due to false positives from very small p-values, likely resulting from uncertainty on the gene-level dispersion parameter estimator in the small sample settings. Additionally shrinking the dispersion estimates strongly increases the performance of the binomial GLM-based workflows and allows for a better type I error rate control. For most datasets, the four different edgeR-based workflows display a similar performance. The two exceptions are panel A1, where the edgeR_optim workflows outper-form the baseline edgeR workflows, and panel D3, where the negative binomial edgeR workflows outperform the quasi-poisson edgeR workflows. As such, the edgeR_NB_optim workflow is the only workflow that performs on par or better than any other workflow across all mock comparisons and performance benchmarks. Hence, we will use this workflow in the case study in Section 2.4.

### 2.3. Combining differential detection and expression results

So far, we have focused solely on DD analyses. However, in real-life applications it will typically be of interest to also assess differential expression (DE) on the same data. Here, we propose a workflow for integrating results from both analyses, which will also be used in Section 2.4 of this manuscript. First, we separately perform a DD analysis on the binarized, pseudobulk aggregated data using the edgeR_NB_optim workflow, and a DE analysis on the pseudobulk aggregated count data using a canonical pseudobulk edgeR analysis. Thus, two p-values will be calculated for each gene. These will be used as input for the stage-wise testing strategy proposed by Van den Berge et al. (2017), which consists of a screening stage and a confirmation stage. In the screening stage, the statistical evidence of the two individual tests are aggregated into an omnibus test. The null hypothesis for the omnibus test is that neither the average detection nor the average expression of the genes is changing between conditions of interest. Genes for which the omnibus null hypothesis is significantly rejected proceed to the confirmation stage, where both hypotheses are assessed separately. The stage-wise testing approach allows us to rank the genes according to their significance on the omnibus test while still providing the detailed resolution on DD and DE for the genes that pass the screening stage. Results can be communicated for both the omnibus test and the individual hypotheses. In Section 4.5, we provide additional background on the stage-wise testing framework.

To assess the performance and type I error control of the stage-wise testing workflow, we re-analyzed the simulated data from Section 2.2. In this simulated data, DD signal between groups was introduced in 5% of the genes by first making gene pairs and then swapping counts between the linked genes in one of the two mock treatment groups. Upon binarization and pseudobulk aggregation of the counts, the genes selected for swapping are assumed to be truly DD between the two treatment groups. We here in parallel perform a DE analysis on the pseudobulk aggregated counts of the same data. Because counts were swapped between genes, the same genes are differentially expressed between groups.

The results of both analyses are displayed in *Supplementary Figure S11, panel A*. The DD and DE workflow are equally powerful in picking up the differential signal. As a consequence, the stage-wise testing analysis does not lead to an increased performance, because the two individual hypotheses provide similar information. The analysis confirms Bouland et al. (2021) findings that a binarized representation of the single-cell expression data allows for a robust description of the relative abundance of transcripts.

In real-life settings, however, we expect more complex differential signals between groups than those generated in our simulation study. Specifically, we also expect some genes to display a pure shift in average expression between groups but with an equal average detection rate, as well as genes with a difference in detection but no shift in the average expression. However, simulating such scenarios while maintaining the original data characteristics is challenging, and there is currently no consensus on which simulator is most suited (Crowell et al., 2022). In addition, simulators typically focus on introducing shifts in average expression and do not allow for introducing shifts in detection. Therefore, we here provide three additional ad-hoc simulation strategies, as described in Section 4.8, to simulate genes that are only DD or only DE. This allows us to conceptually demonstrate the added value of performing a stage-wise analysis.

The results of the performance benchmark on these data are shown in *Supplementary Figure S11, panels B-E*. As expected by design, when we focus the analysis on genes simulated to display a pure DD signal, this signal will only be picked up by the DD analysis, and vice versa (*Supplementary Figure S11, panels B-D*). In these cases, the performance of the omnibus test of the stage-wise analysis workflow is similar to or slightly below the performance of the most relevant analysis, because only one analysis type contains useful information, while the other test statistic reflects noise. However, when analyzing data with all four types of differential signal, the stage-wise testing procedure displays a strong increase in performance over the individual analyses, because there is always at least one of the underlying test statistics that can pick up the differential signal in the omnibus test (*Supplementary Figure S11, panel E*). Note, that the higher performance of the DD analysis compared to the DE analysis is a direct consequence of the design of the benchmark, where more genes were chosen to be DD than DE, and should therefore not be interpreted in terms of the DD signal being easier to detect than DE signal in real-life data.

### 2.4. COVID case study

In this study, we applied our DD workflow to a specific subset of the single-cell RNA-seq dataset obtained from Stephenson et al. (2021). The original publication provided a comprehensive analysis of peripheral blood samples from a cross-sectional patient cohort. The authors employed an approach that integrated single-cell transcriptome analysis, cell-surface protein characterization and lymphocyte antigen receptor repertoire profiling. The cells were stained using the TotalSeq-C antibody cocktail, after which they were loaded on the 10X Chromium platform. Healthy donor samples and samples from COVID patients, ranging from asymptomatic to severely ill patients, were collected across three medical centers in the United Kingdom. For our case study, we focused on five B cell subtypes, for which a sufficient number of cells per patient and a sufficient number of patients per disease status were available. For more details, we refer to Section 4.9.

#### 2.4.1. DE analysis with edgeR

In our study, we first conducted a differential expression analysis using an edgeR-based workflow. This analysis was performed on pseudobulk data, treating each cell type separately. To correct the analysis for sex and sample origin (medical center), we added both variables as fixed effect in the edgeR model. The results of this analysis are summarized in Table 1. The number of DE genes identified between healthy donors and COVID patients are displayed in column 4.

**Table 1:** Number of differentially expressed and differentially detected genes in 25 comparisons between cell types according to COVID status. **Column 1:** an identifier for the comparison (contrast) for easy referencing. Comparisons 18, 20, 23 and 25 are indicated in bold, as these comparisons will be discussed in more detail. **Column 2:** the cell type (B: B cell, CsMB: Class-switched memory B cell, ImmB: immature B cell, NaiveB: naive B cell and UMB: unswitched memory B cell). **Column 3:** the COVID status of the samples that are compared to the healthy samples. **Column 4:** the number of differentially expressed genes identified by the edgeR analysis at the 5% false discovery rate (FDR) level. **Column 5:** the number of differentially detected genes identified by edgeR_NB_optim at the 5% gene-level FDR. **Column 6:** the number of genes that overlap between columns 4 and 5. **Column 7:** the number of genes found in the omnibus test of the stage-wise analysis at the 5% FDR level. **Column 8:** number of differentially detected genes at the 5% gene-level FDR of the stage-wise analysis. **Column 9:** number of differentially expressed genes at the 5% gene-level FDR of the stage-wise analysis.

| Comparison | Cell type | COVID status | DD | DE | Overlap | Omnibus | DD stage2 | DE stage2 |
| --- | --- | --- | --- | --- | --- | --- | --- | --- |
| 1 | B | Asymptomatic | 0 | 0 | 0 | 0 | 0 | 0 |
| 2 | B | Mild | 0 | 2 | 0 | 1 | 0 | 1 |
| 3 | B | Moderate | 7 | 3 | 3 | 5 | 5 | 3 |
| 4 | B | Severe | 1 | 1 | 1 | 1 | 1 | 1 |
| 5 | B | Critical | 0 | 6 | 0 | 0 | 0 | 0 |
| 6 | CsMB | Asymptomatic | 0 | 0 | 0 | 0 | 0 | 0 |
| 7 | CsMB | Mild | 84 | 8 | 6 | 95 | 81 | 33 |
| 8 | CsMB | Moderate | 339 | 191 | 136 | 384 | 326 | 213 |
| 9 | CsMB | Severe | 138 | 50 | 21 | 180 | 138 | 72 |
| 10 | CsMB | Critical | 4 | 11 | 1 | 15 | 6 | 10 |
| 11 | ImmB | Asymptomatic | 1 | 1 | 1 | 1 | 1 | 1 |
| 12 | ImmB | Mild | 48 | 36 | 17 | 71 | 55 | 40 |
| 13 | ImmB | Moderate | 329 | 110 | 93 | 326 | 297 | 154 |
| 14 | ImmB | Severe | 72 | 43 | 29 | 88 | 74 | 52 |
| 15 | ImmB | Critical | 0 | 27 | 0 | 23 | 9 | 19 |
| 16 | NaiveB | Asymptomatic | 0 | 0 | 0 | 0 | 0 | 0 |
| 17 | NaiveB | Mild | 251 | 135 | 101 | 266 | 229 | 161 |
| **18** | NaiveB | Moderate | 2437 | 1417 | 1321 | 2345 | 2239 | 1580 |
| 19 | NaiveB | Severe | 1676 | 558 | 522 | 1529 | 1474 | 812 |
| **20** | NaiveB | Critical | 1307 | 224 | 217 | 1149 | 1125 | 453 |
| 21 | UMB | Asymptomatic | 0 | 0 | 0 | 0 | 0 | 0 |
| 22 | UMB | Mild | 14 | 19 | 3 | 36 | 21 | 21 |
| **23** | UMB | Moderate | 119 | 164 | 60 | 232 | 159 | 162 |
| 24 | UMB | Severe | 78 | 11 | 6 | 99 | 84 | 30 |
| **25** | UMB | Critical | 0 | 50 | 0 | 16 | 3 | 14 |

#### 2.4.2. DD analysis

Following the DE, we proceeded to conduct a DD analysis separately for each cell type, again including a fixed effect for sex and medical center. For this analysis, we used the top-performing method from our simulation study, edgeR_NB_optim. In Table 1, we present the results of this DD analysis, specifically highlighting the number of DD genes for each contrast involving patients with a particular COVID disease status vs healthy patients in column 5. From Table 1, it is evident that asymptomatic COVID patients, regardless of cell type, exhibit no DD nor DE genes compared to healthy patients at the 5% FDR level. However, for comparisons involving more severe disease statuses a substantial number of DE an DD genes does not overlap for particular cell types, highlighting the complementary nature of our DD analysis in relation to the outcomes of the conventional DE analysis. Moreover, the number of significant differential genes is larger for the DD analysis than for the DE analysis in several comparisons.

To assess whether the DD analysis provides additional biological insights compared to the DE analysis alone, we conducted a focused investigation on four distinct cell type-COVID status combinations: Naive B cells in moderately and critically ill patients, as well as unswitched memory B cells in moderately and critically diseased patients (comparisons 18, 20, 23 and 25, respectively). The naïve B cells were selected because they had the highest number of differential genes. The unswitched memory B cells comparisons between healthy donors and moderately and critically ill patients are comparisons where more differential signal was picked up in the DD analysis than in the DE analysis. For each comparison, we identified and examined the top 10 DD genes that were not picked up as DE on the 5% false discovery rate level, evaluating their relevance to COVID (Supplementary Table 1).

For this subset of genes, several genes are already known to be associated with COVID. We here enumerate the genes within our subset of genes that have been identified previously as associated with COVID. ATP5F1A (Guarnieri et al., 2022), IGHV6-1 (Yang et al., 2022), ACTG1 (Yuka and Yilmaz, 2021), GDI2 (Lata et al., 2022), IGLV1-51 (Zhao et al., 2021) and PKM (Qi et al., 2021)) for the naïve B cells in moderately ill patients; IFITM1 (Arefinia et al., 2023), IGHV6-1 (Yang et al., 2022) and S100A8 (Mellett and Khader, 2022) for the naive B cells in critically ill patients; PSME2 (Alfaro et al., 2022), IGKC (Momeni et al., 2023), PMAIP1 (Park and Lee, 2020), DDX21 (Henriques-Pons et al., 2022), EIF2AK2 (Melnichuk et al., 2022) and CD27 (Wen et al., 2020) for the unswitched memory B cells in moderately ill patients; and DDX21 (Henriques-Pons et al., 2022), IGKC (Momeni et al., 2023), COX6A1 (Mukund et al., 2021), CD27 (Wen et al., 2020) and RHOA (Khatoon et al., 2020) for the unswitched memory B cells in critically ill patients. The significant number of genes with a potential link to the disease highlights the valuable biological insights gained from a DD analysis, beyond those obtained with a conventional DE analysis.

Figure 3 shows the added value of combining information from DE and DD analyses. In panel A, we plot the p-values from the DD analysis against the p-values of the DE analysis (upon -log10 transformation) for the comparison 18 of Table 1. While many genes are both significantly DD and DE (green color), many genes are only DE (red color) or only DD (pink color). We highlight three genes in panel A, TRBC2, PPIB and ATP5F1A, which we will consider archetypes of genes that are both DE and DD, only DE or only DD, respectively. In panels B-D of Figure 3, the genes highlighted in panel A will be inspected in detail.

**Figure 3:**
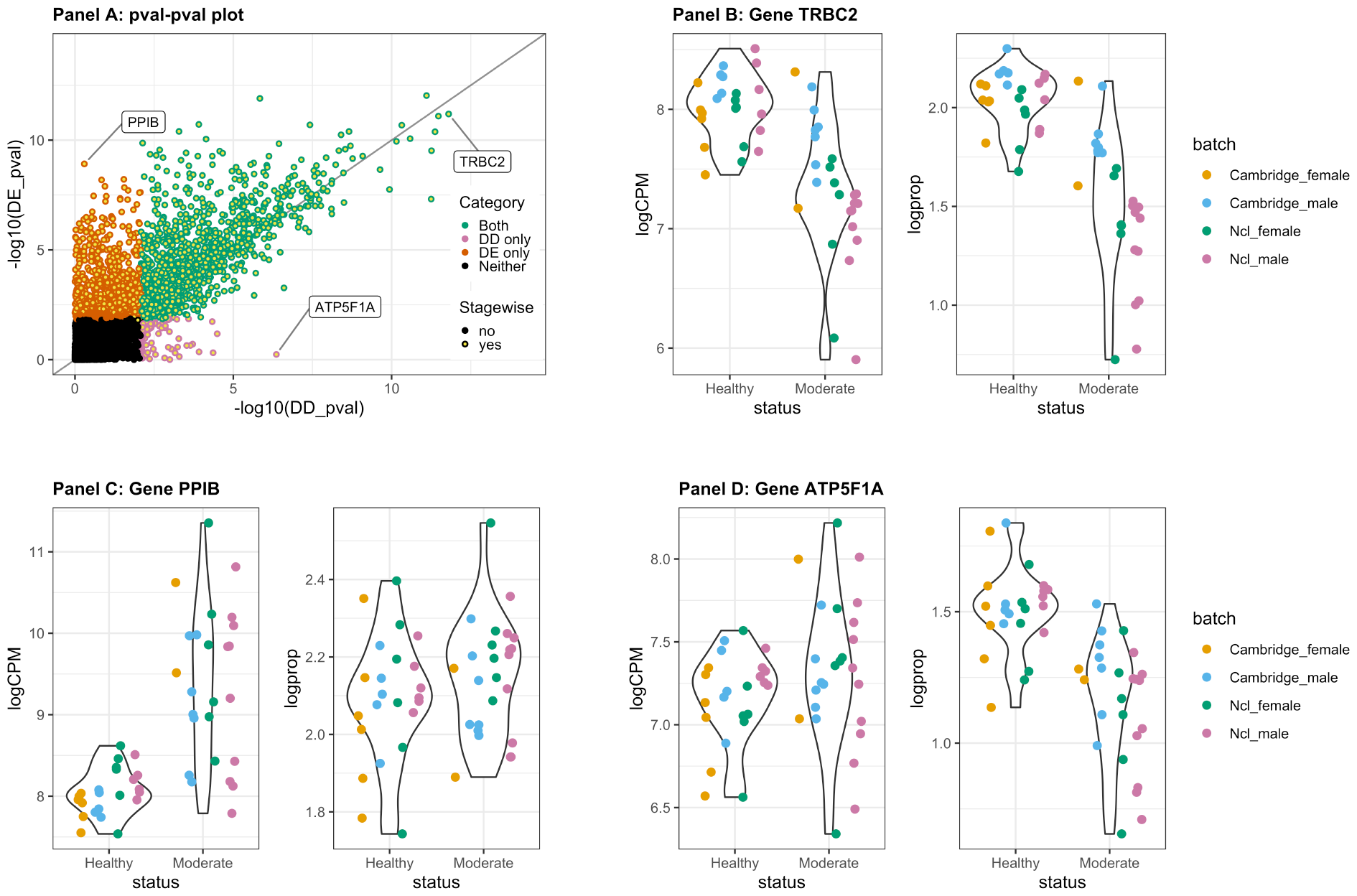
Added value of combining information from DE and DD analyses. This figures focuses on comparison 18 from Table 1. **Panel A:** DE and DD analysis transformed p-values. The p-values from the DD analysis are displayed against the p-values of the DE analysis (upon -log10 transformation). Each dot represents the statistical significance of a single gene in the DE and DD analysis. Statistical significance was assessed at an FDR of 5%. Each is either both DE and DD (green), only DE (red), only DD (pink) or neither DE nor DD (black). Genes that passed the screening stage of the stage-wise testing procedure are indicated with a yellow center in the dot. Three genes are highlighted in this panel: TRBC2, PPIB and ATP5F1A. These will be considered archetypes for genes that are both DE and DD, only DE or only DD, respectively. **Panel B:** Violin plots for gene TRBC2. The left panel displays a measure of gene expression, while the right panel displays a measure for gene detection. In line with panel A, this gene has a clear signal of both DE and DD. **Panel C:** Violin plots for gene PPIB. In line with panel A, this gene has a strong DE signal, but no evidence for DD. **Panel D:** Violin plots for gene ATP5F1A. In line with panel A, this gene has a strong DD signal, but no evidence for DE.

In Figure 3 panel B, we visualize the expression and detection of gene TRBC2 (T-cell receptor *β* constant 2) using violin plots, stratified between healthy donors and moderately ill COVID patients. The sample-specific expression and detection values are overlaying the violin plots, with the coloring indicating the batch effect that was used as a fixed effect in our analyses (see Section 4.9). On the left, the expression is shown in terms of log2-transformed counts-per-million (logCPM). The average logCPM expression is higher for healthy donors than for moderately ill patients (FDR = 9.46e-09). On the right, detection is visualized in terms of the normalized, log2-transformed proportion of detection (logprop). The interpretation of this measure is as follows: a value of 1 indicates that the detection of the gene in the sample is twice as high as the average detection across all genes in that sample. For gene TRBC2, the average logprop is higher for healthy donors than for moderately ill patients (FDR = 9.80e-09). Panel C visualizes the expression and detection of gene PPIB (Peptidylprolyl Isomerase B), which is a gene known to be involved in COVID viral cell-entry (Bergsneider et al., 2021). PPIB displays a clear DE signal (FDR = 2.23e-07), but no statistical evidence for DD was observed (FDR = 0.70). Finally, panel D is visualizing gene ATP5F1A. For this gene, no statistical evidence of DE (FDR = 0.75) and strong evidence for DD (FDR = 4.04e-05) was obtained. ATP5F1A (ATP Synthase F1 Subunit Alpha) is a subunit of the mitochondrial ATP Synthase protein complex, which is crucial in the process of oxidative phosphorylation (Guarnieri et al., 2022). Chen et al. (2023) previously discussed how the lack of components of the ATP Synthase protein complex may cause defects in the complex, jeopardizing its functionality. While the overall expression of ATP5F1A across all naive B-cells is equal between healthy donors and moderately ill patients, the lower average detection in the individual cells of the latter indicates a lower proportion of individual cells in which ATP5F1A is present. As such, there may be fewer cells in moderately ill patients that are capable of producing a functional ATP Synthase protein complex.

Note that visualizations for the other three comparisons (comparisons 20, 23 and 25) are available from *Supplementary Figures S12-S14*. For comparison 25 only panel A is shown, because no significant DE features were identified in this comparison.

#### 2.4.3. Stage-wise analysis

To integrate information from the DD and DE analyses, a stage-wise analysis was performed. The results of the omnibus test, displayed in column 7 of Table 1, reveal the number of genes that pass the screening stage for each cell type and COVID status. With the screening stage, we refer to gene-level omnibus tests that are conducted based on aggregated evidence across both the DD and DE analysis results. It is important to note that the stage-wise analysis often uncovers more results than the sum of the two separate analyses (upon collapsing genes identified by both analyses). In Figure 3, genes passing the screening stage are indicated with a yellow center in the dot. Several genes that pass the 5% FDR significance threshold for the individual DE or DD hypothesis do not pass the screening stage (full red or pink dots). When there is only differential signal in one of the two analyses, this signal is to some extent diluted in the p-value aggregation step of the stage-wise testing procedure, leading to a potential loss of the signal. Conversely, some genes that do not pass the 5% FDR significance threshold for the individual DE and DD hypothesis are being picked up in the stage-wise testing approach (black dots with yellow centers, see *Supplementary Figures S12-S14* for examples). This scenario arises when the individual hypotheses both display some evidence for differential signal, but when this signal is insufficient for the individual hypotheses to be rejected.

For the genes that pass the screening stage, the stage-wise testing procedure allows for performing post-hoc tests to pinpoint if these genes are DD, DE or both. We refer to these results as the stage2 DD and stage2 DE results. The number of stage2 DD and stage2 DE genes for each comparison in our case study are displayed in Table 1, columns 8 and 9, respectively. These numbers are highly similar to those of the individual DD and DE analyses from Table 1, columns 4 and 5, respectively. In Supplementary Table 2, we additionally show that the genes uncovered by the individual analyses and those uncovered in the second stage overlap strongly. Hence, we generally recommend to use the two-stage testing framework, as (1) it allows for interpreting the results both in terms of the omnibus hypothesis as the individual DD and DE hypotheses and (2) it ensures that the gene-level FDR is controlled at the significance level *α*. These recommendations will be discussed more thoroughly in Section 3.

**Table 2:** GSEA Results for Unswitched Memory B Cells of Moderately Ill Patients. **Column 1:** The gene ontology (GO) terms of the processes that are significantly enriched. **Column 2:** Ranking of the GO terms from column 1 when performing a GSEA on the significant genes from the DD analysis (stage2). **Column 3:** Ranking of the GO terms from column 1 when performing a GSEA on the significant genes from the DE analysis (stage2). **Column 4:** Ranking of the GO terms from column 1 when performing a GSEA on the significant genes from the stage-wise analysis, in which the results from the DD and DE analyses are aggregated. **Column 5:** P-values for the GO terms in column 1 when performing a GSEA on the significant genes from the DD analysis. **Column 6:** P-values for the GO terms in column 1 when performing a GSEA on the significant genes from the DE analysis. **Column 7:** P-values for the GO terms in column 1 when performing a GSEA on the significant genes from the omnibus test of the stage-wise analysis.

| GO Term | Rank<br>DD<br>stage2 | Rank<br>DE<br>stage2 | Rank<br>Omni-<br>bus | pvalue<br>DD<br>stage2 | pvalue<br>DE<br>stage2 | pvalue<br>Omni-<br>bus |
| --- | --- | --- | --- | --- | --- | --- |
| Response to virus | 1 | 19 | 12 | 4.1e-13 | 1.3e-08 | 1.3e-10 |
| Immune response | 2 | 13 | 10 | 8.0e-12 | 6.1e-10 | 3.5e-11 |
| Defense response to symbiont | 3 | 23 | 16 | 1.9e-11 | 6.0e-07 | 2.0e-09 |
| Defense response to virus | 4 | 24 | 17 | 1.9e-11 | 6.0e-07 | 2.0e-09 |
| Response to stress | 5 | 56 | 32 | 1.1e-10 | 1.4e-04 | 1.8e-07 |
| Immune system process | 6 | 21 | 21 | 1.6e-10 | 1.3e-07 | 7.2e-09 |
| Response to other organism | 7 | 27 | 23 | 5.4e-10 | 2.7e-06 | 1.4e-08 |
| Response to external biotic stimulus | 8 | 28 | 24 | 5.7e-10 | 2.9e-06 | 1.5e-08 |
| Response to interferon-alpha | 9 | 32 | 20 | 8.1e-10 | 5.1e-06 | 6.4e-09 |
| Response to biotic stimulus | 10 | 30 | 27 | 1.1e-09 | 4.6e-06 | 3.0e-08 |
| Cytoplasmic translation | 1281 | 1 | 1 | 2.5e-01 | 3.7e-29 | 1.8e-25 |
| Translation | 748 | 2 | 2 | 1.0e-01 | 1.3e-23 | 5.2e-19 |
| Peptide biosynthetic process | 824 | 3 | 3 | 1.2e-01 | 4.0e-23 | 1.6e-18 |
| Amide biosynthetic process | 1131 | 4 | 4 | 2.0e-01 | 5.9e-21 | 1.8e-16 |
| Peptide metabolic process | 1246 | 5 | 5 | 2.4e-01 | 3.5e-20 | 1.0e-15 |
| Cellular macromolecule biosynthetic process | 1404 | 6 | 6 | 3.0e-01 | 1.1e-17 | 3.0e-13 |
| Cellular amide biosynthetic process | 1717 | 7 | 8 | 4.7e-01 | 1.1e-16 | 1.9e-12 |
| Organonitrogen compound biosynthetic process | 1345 | 8 | 13 | 2.8e-01 | 2.4e-14 | 1.4e-10 |
| Gene expression | 51 | 9 | 7 | 3.4e-04 | 3.2e-14 | 7.3e-13 |
| Cellular nitrogen compound biosynthetic process | 54 | 10 | 9 | 3.9e-04 | 1.5e-12 | 7.8e-12 |

#### 2.4.4. Gene set enrichment analysis

We again focus on comparisons 18, 20, 23 and 25 from Table 1 to evaluate the use of a stage-wise workflow in the functional interpretation of biological experiments. More specifically, we have performed three gene set enrichment analysis (GSEA) runs, using as input the significant genes obtained from (1) the stage2 DD result, (2) the stage2 DE result, and (3) the results from the omnibus test of the stage-wise testing procedure. The GSEA was performed using the *goana* function from the R package limma (Smyth, 2004). For each GSEA run, we retained the top 10 significantly enriched gene ontology (GO) terms. In Table 2, we display the GSEA results for comparison 23. In column 1 of Table 2, we display the union of the top 10 GO terms across the three GSEA runs. In columns 2, 3 and 4, we show the ranking of the GO terms in the three respective GSEA runs (stage2 DD, stage2 DE and omnibus). In columns 5, 6 and 7, we show the p-value for the enrichment of the GO terms in the three respective GSEA runs.

First, we note that the GO terms obtained from the stage2 DD results are different from those obtained from the stage2 DE results. None of the top 10 ranking GO terms from the stage2 DD analysis overlap with the top 10 GO terms from the stage2 DE analysis, and vice versa. Moreover, most of the top GO terms from the DE analysis are not picked up as significant by the GSEA run on the DD results. This demonstrates that the DD and DE analyses are providing complementary information also on the functional level.

Secondly, the GO terms from the stage2 DD analysis are all pointing towards a immune response to a viral infection. The GO terms from the stage2 DE analysis are instead more generic terms, some of which have been reported previously to be enriched in COVID patients (e.g., peptide biosynthetic process (Huang et al., 2023), organonitrogen compound biosynthtic process (Jeyananthan, 2022), gene expression (Choudhary et al., 2021), cellular amide metabolic process (Huang et al., 2023), amide biosynthetic process (Li et al., 2021) and cellular macromolecule biosynthetic process (Huang et al., 2023)).

The results for the other comparisons (18, 20 and 25) are provided in *Supplementary Tables S3-S5*. The complementarity between the stage2 DD and stage2 DE GSEA runs is observed in all comparisons. However, many of the reported GO terms for both runs are generic terms. As such, it is not clear if either of the two GSEA runs provides more biologically relevant information than the other. Note that for comparison 25 we only show the GSEA run on the stage2 DD results, as no DE genes were obtained in this comparison.

Finally, the ranking of GO terms and the corresponding p-values are similar between the stage2 DE results and the omnibus results, for all comparisons (Table 2 and *Supplementary Tables S4 and S5*). In addition, the ranking of the GO terms and corresponding p-values are also similar between the individual DE results and the stage2 DE results, and between the individual DD results and the stage2 DD results *(Supplementary Tables S6-S9*). Altogether, this indicates that the GSEA results for the omnibus, DE and DE stage2 results are similar. However, using the stage-wise testing framework has the additional advantage of being able to assess the individual stage2 DE and DD results in a unified framework that correctly controls the type I error at the gene level.

### 2.5. Lupus case study

We applied our DD workflow to a specific subset of the single-cell RNA-seq dataset obtained from (Perez et al., 2022). The original publication utilized mux-seq technology (Rashmi et al., 2022) to profile over 1.2 million peripheral blood mononuclear cells derived from 162 systemic lupus erythematosus (SLE) cases and 99 healthy controls of Asian or European ancestry. For our case study, we perform a DE and DD analysis between 22 healthy and 26 ill European women for three cell types (see Section 4.10).

As for the previous case study, a DE analysis was conducted with edgeR. The number of DE genes with an FDR below 5% is shown for each cell type in column 1 of Table 3. Next, we conducted a DD analysis separately for each cell type using edgeR_NB_optim. We first visualize the p-values of the DE analysis against the p-values of the DD analysis (*Supplementary Figure S15*). Compared to the corresponding figures for the COVID case study (Figure 3 and *Supplementary Figures S12-S14*), the DE and DD p-values in the three cell types of the lupus dataset are more concordant, which suggests that the two analyses are providing similar information. Next, we report the number of significant genes at the 5% FDR in the individual DE and DD analysis, as well as the number genes that pass the omnibus test of the stage-wise testing procedure (Table 3, columns 2, 3 and 4, respectively). In line with the results from our COVID case study, the omnibus test yields a higher number of significant genes than the two individual analyses. The number of DD and DE genes that are identified in the second stage of the stage-wise testing procedure are shown in columns 5 and 6.

**Table 3:** Number of differentially expressed and differentially detected genes in 3 different cell types. **Column 1:** cell type. **Column 2:** the number of significantly differentially expressed genes identified by the edgeR analysis (5% FDR). **Column 3:** the number of significantly differentially detected genes identified by edgeR_NB_optim (5% FDR). **Column 4:** the number of genes that pass the omnibus test of the stage-wise testing procedure (5% FDR). **Column 5:** the number of significantly differentially expressed genes in the second stage of the stage-wise testing procedure. **Column 6:** the number of significantly differentially detected genes in the second stage of the stage-wise testing procedure. **Column 7:** the number of overlapping genes between the individual DE and DD analysis results. **Column 8:** the number of overlapping genes between the stage two DE analysis and the stage two DD analysis results. **Column 9:** the number of overlapping genes between the individual DE analysis and the stage two DE analysis results. **Column 10:** the number of overlapping genes between the individual DD analysis and the stage two DD analysis results.

| Cell Type | DE | DD | Omnibus | DE stage2 | DD stage2 | Overlap DE-DD | Overlap DE stage2 - DD stage2 | Overlap DE - DE stage2 | Overlap DD - DD stage2 |
| --- | --- | --- | --- | --- | --- | --- | --- | --- | --- |
| T4 Naive | 248 | 170 | 264 | 242 | 189 | 146 | 167 | 237 | 170 |
| B memory | 58 | 33 | 64 | 56 | 36 | 27 | 28 | 54 | 33 |
| ncM | 340 | 266 | 382 | 334 | 277 | 209 | 229 | 329 | 259 |

As in the COVID case study, these numbers are similar to the numbers obtained by the individual DE and DD analysis from columns 2 and 3. We note that while the DE and DD p-values show a strong concordance (*Supplementary Figure S15*), the set of DE genes and DD genes does not overlap fully (Table 3, columns 7 and 8). As such, there is still added value in performing both analyses. Finally, the genes identified in the individual DE and DD analyses nearly perfectly overlap with those identified in the DE and DD analyses from the second stage of the stage-wise testing procedure (columns 9 and 10). As such, even though the DE and DD p-values show a strong concordance (*Supplementary Figure S15*), we observe no disadvantage of performing a stage-wise testing analysis.

## 3. Discussion

In this manuscript, we have presented a workflow for performing differential detection (DD) analyses on scRNA-seq data.

First, we have benchmarked eight DD workflows that were either based on a binomial GLM or on edgeR. For the binomial models, we showed that working under a quasi-likelihood framework improved the type I error control, while including a normalization offset and stabilizing the estimation of the quasi-binomial overdispersion parameter increased statistical power. The edgeR-based models also properly control the type I error and displayed a higher power than the binomial models. By modeling the data at the pseudobulk level, we correctly address the within-sample correlation and reduce the computational complexity.

Next, we have shown how results from a DD analysis and a classical differential expression (DE) analysis can be integrated. In two case studies, we show how combining DD an DE results allows for uncovering biologically relevant signal that could not be detected with a classical DE analysis alone. Furthermore, the resolution of the hypothesis that can be tested is increased, as our workflow can pinpoint if the differential signal is due to differences in detection, differences in average expression, or both.

We analyzed data originating from droplet-based scRNA-seq platforms, i.e., 10X Chromium and mux-seq (Rashmi et al., 2022) for the COVID and Lupus datasets, respectively. While these platforms allow for characterizing several thousands of cells per sample, the resulting gene expression count matrix is notoriously sparse. For the processed single-cell level data in the COVID case study, after gene-level and cell-level filtering, 86% of the counts are zeros, 9% are ones, 2% are twos and 3% are counts larger than 2. Similarly, for the lupus dataset, the final single-cell count matrix consisted of 84% zeros, 10% ones, 2% twos and 4% counts larger than two. Hence, analyzing the data at the single-cell level using Gaussian models on log-transformed counts as proposed by MAST (Finak et al., 2015) can be expected to be problematic, as discussed by Townes et al. (2019).

We have proposed performing DD analyses and DE analyses separately, after which the results from the two analyses are integrated using the two-stage testing paradigm proposed by Van den Berge et al. (2017). Alternatively, the inference on the detection component and the count component could have been obtained by using a hurdle model, as implemented in MAST (Finak et al., 2015). A hurdle model has two components: a Bernoulli component modeling presence/absence (i.e., detection), and a component that models the positive counts to infer on DE given that the gene is expressed in a cell. Hurdle models imply an estimation orthogonality, which allows the parameters of each component to be estimated separately by only considering a binary indicator for presence/absence for estimating the parameters to infer DD and the positive counts for estimating the parameters to infer DE (e.g. (Finak et al., 2015)). Conceptually, the main difference between our workflow and a hurdle model is in the modeling of the count component. Hurdle models use a zero-truncated model for the counts (i.e., a zero-truncated Gaussian model in the case of MAST), whereas we use a full count distribution. However, because we aggregate the data on the pseudobulk level, our data will be much less sparse than regular single-cell data (e.g., 2.4% zeros in the COVID case study data and 1.4% zeros in the lupus case study data). Hence, we do not expect a large impact of truncating the count model.

In our case studies, we have demonstrated that DE and DD analysis results provide complementary information. One open question is the underlying mechanism that is driving this observed complementarity. It is possible that it is a consequence of true differences in the underlying biology, where cells truly display DE for some genes and DD for other genes. An alternative explanation is that the underlying biological mechanism is introducing DE in reality, but that for lowly expressed genes we gain power by testing for DD.

As a guideline for differential RNA-seq data analysis, we recommend performing both a DE and a DD analysis on pseudobulk RNA-seq data. For the DD analysis, we recommend using our edgeR_NB_optim workflow, which is highly scalable, provides good type I error control and generally has a higher power than the other methods in our benchmark. Next, we recommend using the stage-wise testing framework from (Van den Berge et al., 2017). This allows for testing both the omnibus null hypothesis that a gene is neither DD nor DE, as well as performing the individual DD and DE tests while correctly controlling the gene-level FDR. As demonstrated in our case studies, the DE and DD analysis provided complementary information, emphasizing the importance of modeling both components.

## 4. Methods

### 4.1. Differential detection analysis for scRNA-seq data

The first step of our proposed differential detection analysis workflow is to binarize the input scRNA-seq gene expression matrix by setting every non-zero expression value to a value of one. The binarized single-cell matrix could in principle be used directly for DD inference. However, when cells from the same sample (e.g. subject) are being profiled, the cell-level expression profiles are no longer independent, and this within-sample correlation should be accounted for in the statistical inference (Lun and Marioni, 2017, Zimmerman et al. (2021) and Squair et al. (2021)). We therefore adopt a similar count aggregation strategy on the binarized counts as proposed by Crowell et al. (2020) for conventional scRNA-seq counts, i.e., taking the sum of the sample-level binarized counts separately for each cell type, thus creating sample-level pseudobulk binomial counts. Because the summation is performed on binarized data, the resulting counts have the interpretation of the number of cells in which a gene is expressed in each sample and cell type combination. Finally, a DD analysis is performed for each cell type separately. In our mock comparison and performance evaluation benchmarks, DD models for both binarized single-cell data and pseudobulk binomial data will be evaluated.

### 4.2. Modeling differential detection with binomial generalized linear models

We benchmark four generalized linear modeling (GLM) strategies from the binomial family to perform differential detection analyses on pseudobulk binomial data. In order of increasing complexity, these four models are

- a binomial GLM
- a quasi-binomial GLM
- a quasi-binomial GLM with a normalization offset
- a quasi-binomial GLM with a normalization offset and shrinkage of the dispersion parameter.

In this paragraph, we use the following notation. Consider a dataset with *G* genes and *N* samples. For each gene *g* = 1, …, *G*, we model the number of times the gene was detected in sample *i* = 1, …, *N*, a quantity that we represent with the random variable *Y*_*gi*_. For pseudobulk data, we assume *Y*_*gi*_ to follow a Binomial distribution *Y*_*gi*_ = *Bin*(*π*_*gi*_, *n*_*i*_), with *π*_*gi*_ the expected probability of detecting gene *g* in sample *i* (successes), and *n*_*i*_ the number of cells that were aggregated for sample *i* (trials). For single-cell level data, *n*_*i*_ = 1, this reduces to a Bernoulli distribution. Note that for each modeling strategy, we fit these models separately for each cell type.

#### 4.2.1. Binomial GLM

Using binomial regression, *Y*_*gi*_ can be modeled as follows:

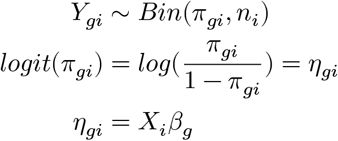

We model *π*_*gi*_ using a logit link function, where *β*_*g*_ is a *p* × 1 column vector of regression parameters associating the logit average detection with the experimental covariates. *β*_*g*_ is estimated using maximum likelihood. Finally, *X*_*i*_ is a row in the *n* × *p* design matrix *X* that corresponds with the covariate pattern of sample *i*, with *p* the number of parameters of the mean model, i.e., the length of vector *β*_*g*_.

The variance of the binomial model for gene *g* is given by

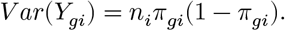

For each gene *g*, we may then consider the following null and alternative hypotheses:

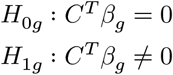

with *C*^*T*^ a contrast 1 × *p* vector that allows for testing linear combinations of *β*_*g*_. Inference is done with a Wald test, which is assumed to asymptotically follow a *z*-distribution under the null hypothesis of no association.

#### 4.2.2. Quasi-binomial GLM

The simple binomial regression model imposes a strong assumption on the mean-variance relationship in the data. To allow for more flexibility, we may use a quasi-binomial GLM. The quasi-binomial GLM models the first two moments (the mean and variance) according to the following expressions:

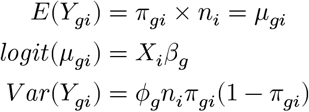

with *μ*_*gi*_ the unobserved average expression of gene *g* in pseudobulk sample *i*. Compared to the canonical binomial variance function described in the previous paragraph, the variance function of the quasi-binomial GLM has a dispersion parameter *ϕ*_*g*_. This dispersion parameter allows for describing over- or underdispersion of the data with respect to the canonical binomial variance. The estimation equations for the mean model parameters of the binomial and quasi-binomial models are equal. Both approaches only differ in their variance estimator on the mean model parameters. Indeed, the quasi-binomial model simply rescales the variance estimator of the binomial with the dispersion parameter. Inference is done using approximate t-tests that follow a t-distribution with *n* − *p* degrees of freedom under the null hypothesis.

#### 4.2.3. Quasi-binomial GLM with normalization offset

The binomial GLM and quasi-binomial GLM described above are modeling the association between the average detection of gene *g* and the experimental covariates. The linear predictor can be interpreted as the log-odds of detecting gene *g* given a covariate pattern. Contrasts of the model parameters thus can be interpreted as log-odds ratios, e.g. the ratio of the log-odds of detecting gene *g* in one covariate pattern versus another covariate pattern.

However, note that there can be systematic differences in the overall detection rate across all genes between the different samples or between the individual cells that may relate to technical variation (Hicks et al., 2018). For instance, when there are systematic differences in sequencing depth between batches, it can be expected that the average detection of all genes (i.e., the fraction of genes detected in a sample) is higher in the batch that was sequenced more deeply. To account for differences in the overall detection across all genes between the different samples or individual cells, we include an offset in the quasi-binomial model, such that

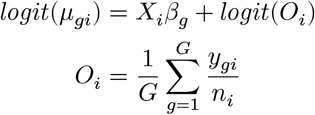

with *O*_*i*_ the sample- or cell-specific offset for the average detection over all genes. By including *O*_*i*_ as an offset to the quasi-binomial GLM, the contrast *X*_*i*_*β*_*g*_ can be interpreted as the log-odds ratio of detecting gene *g* given covariate pattern *X*_*i*_ versus the average detection over all genes in the same sample or cell. This normalization offset corresponds to the cellular detection rate that is used by MAST (Finak et al., 2015).

#### 4.2.4. Quasi-binomial GLM with normalization offset and shrinkage

As described above, the quasi-binomial model estimates a dispersion parameter *ϕ*_*g*_ to account for over- or underdispersion in the data with respect to the binomial variance. In bulk RNA-seq data, the dispersion parameter *ϕ*_*g*_ is typically highly variable between genes and the number of observations is often small, resulting in imprecise estimates for *ϕ*_*g*_. While single-cell data has thousands or even millions of observations, this number again is reduced when performing pseudobulk aggregation. As such, we adopt the strategy proposed for bulk RNA-seq data (Smyth, 2004, and Lund et al. (2012)), which stabilizes the estimation of the dispersion parameter by borrowing strength across genes and shrinking the dispersion estimates using empirical Bayes. This procedure is implemented in the *squeezeVar* function of the *limma* Bioconductor package. After thus having obtained a shrunken dispersion estimate 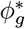, inference is achieved by computing a moderated t-statistic as described in Smyth (2004).

### 4.3. Modeling differential detection with edgeR

We assess four differential detection workflows that use a quasi-poisson or quasi-negative binomial approximation to the binomial model. For this, we build on the popular DE tool edgeR (Robinson et al., 2010, and Lund et al. (2012)). First, we again binarize and perform pseudobulk aggregation on the original gene expression abundances. Next, we utilized a quasi-negative binomial model by employing the *glmQLFit* function from the *edgeR* package to perform a DE analysis on each cell type separately. Like the quasi-binomial GLM workflows described above, edgeR accounts for gene-specific variability by estimating an additional over-dispersion parameter. The quasi-negative binomial model considers the number of times gene *g* is detected in sample *i* (denoted as *Y*_*gi*_). *Y*_*gi*_ is modeled using the following quasi-negative binomial model:

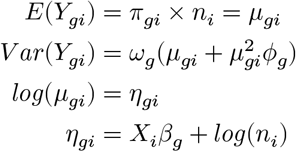

where *π*_*gi*_ represents the expected count for gene *g* in sample *i* given the number of cells *n*_*i*_ and covariate pattern *X*_*i*_ for sample *i, μ*_*gi*_ the unobserved average expression of gene *g* in pseudobulk sample *i*,and *ϕ*_*g*_ is the NB dispersion parameter and *ω*_*g*_ is the quasi-likelihood dispersion parameter. To account for the total number of aggregated cells in pseudobulk sample *i, n*_*i*_ is included as an offset term. Finally, *β*_*g*_ is a vector of regression parameters modeling the association between the average detection and the experimental covariates.

Furthermore, we also employed a quasi-poisson model by setting the dispersion parameter *ϕ*_*g*_ to zero. The quasi-poisson model is specified as follows:

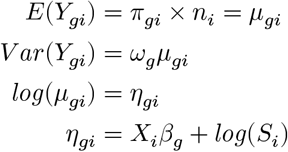

where again *μ*_*gi*_ represents the expected count for gene *g* in sample *i* given the number of cells *S*_*i*_ and covariate pattern *X*_*i*_, and *ω*_*g*_ is the quasi-likelihood dispersion parameter.

We have also introduced two optimized versions of the *edgeR* workflow, which we named edgeR_NB_optim and edgeR_QP_optim. In this workflow, we remove genes that are detected in more than 90% of the cells across all samples. The rationale behind this filtering is that genes that are nearly always expressed in nearly all samples exhibit minimal differences between groups, leading to p-values peaking at 1. Furthermore, the dispersion estimate for such genes is typically artificially low. As edgeR performs shrinkage of the dispersion estimates across genes, these artificially low estimates could have a strong impact on the estimates of the other genes. Across our different benchmarks and case studies, this additional gene-level filtering removed between 1% and 6% of the genes. In addition, edgeR does not allow the variance to be below the Poisson variance by default. For edgeR_NB_optim and edgeR_QP_optim, we also allow for a variance below the Poisson variance by setting the *poisson*.*bound* argument of the *glmQLFTest* of edgeR to false. Finally, we again include the cellular detection rate as an additional model offset as described in Section 4.2.3. The resulting model is thus specified as follows:

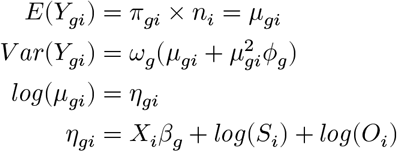

where all features are the same as those described for the quasi-negative binomial and *O*_*i*_ represents the cellular detection rate.

### 4.4. Mock simulations

The type I error of the different methods was benchmarked in two simulation studies. The simulated data is based on two real datasets: the first is a study on systemic lupus erythematosus (Perez et al., 2022), the second is a large COVID dataset (Stephenson et al., 2021).

For the simulations based on the lupus dataset (Perez et al., 2022), we filtered the data to obtain a homogeneous subset retaining only samples from healthy, European women under 50 years old. Additionally, only subjects from three specific sequencing batches were included (batches “dmx_count_AH7TNHDMXX_YE_8-30”, “dmx_count_AHCM2CDMXX_YE_0831” and “dmx_count_BH7YT2DMXX_YE_0907”), thus retaining 44 patients. To further enhance homogeneity, we analyzed cell types separately, focusing on T4 naive cells, B memory cells, and non-classical myeloid cells. These three cell types were selected because they were observed in all patients, and because they have a high, intermediate and relatively low number of cells per patient, respectively. The median number of cells per patient is 708 cells for the T4 naive cell type, 321 cells for the B memory cell type and 118.5 cells for the non-classical myeloid cells. Gene-level filtering was performed separately for each cell type, only retaining genes that have an expression of at least 10 unique molecular identifyers (UMIs) over all cells, and are expressed in at least 2% of all cells. The 44 patients were randomized into two mock groups, so none of the genes were expected to be DD or DE between the mock groups. We repeated this random assignment five times.

To assess the impact of sample size on performance, we additionally downsampled the dataset for each cell type to perform mock analyses on 5 vs 5, 10 vs 10, and 15 vs 15 patients. For each sample size, the subsampling was performed five times, each time with a different randomly selected subset of samples.

For the COVID dataset, we retained patients from the Cambridge and Newcastle sequencing sites. Only individuals classified as healthy, mildly ill, moderately ill, critically ill or severely ill were considered, as the other groups did not contain sufficient patients. We again focused on three cell types: naive B cells, class-switched memory B cells and immature B cells. These three cell types were selected because they have a high, intermediate and relatively low number of cells per patient. Naive B cells were detected in 99 patients, with a median of 256 cells per patient. Class-switched memory B cells were detected in 76 patients, with a median of 54.5 cells per patient. Finally, immature B cells were detected in 69 patients, with a median of 38 cells per patient. Mock analyses and simulations were conducted separately for each cell type and disease status combination. Furthermore, when assigning patients to mock groups, patients were balanced for gender and sequencing site to account for possible confounders. Again, the subsampling was performed five times, each time with a different randomly selected subset of samples.

### 4.5. Stage-wise testing

We integrate the results from the DD analysis and DE analysis using the two-stage testing framework from Van den Berge et al. (2017).

In the first stage, we identify differential genes by using an omnibus test for DD and DE. The null hypothesis for the omnibus test is that neither the average detection nor the average expression of the genes is changing between conditions of interest. To construct the omnibus test, we must aggregate the statistical evidence for the DD and DE tests. We adopt the strategy proposed by Wilson (2019) that can aggregate p-values that are correlated. Benjamini-Hochberg FDR multiple testing correction (Benjamini and Hochberg, 1995) is performed on the aggregated p-values, and all genes with an FDR-adjusted aggregated p-value below the significance level *α* are selected to proceed to the second stage.

In the second stage, we perform post-hoc tests on the differential genes from stage one to unravel whether they are DD, DE or both. If differential signal is present in both individual tests, the stage-wise testing framework has increased power for detecting this signal. If the signal is present in only one of the two tests, then the power of stage-wise testing framework will be lower than that of the individual hypotheses, due to a dilution of the signal. Note that this stage-wise testing framework ensures that the gene-level FDR is controlled at the significance level *α*.

### 4.6. Simulations including evidence for DD and DE

To generate simulated datasets with differentially expressed genes, we employed the following approach; after having assigned patients to the two mock groups as previously described, we randomly swapped counts between 5% of the genes in one of the mock treatment groups. This swapping of gene counts introduces DD and DE of different magnitudes between the mock treatment groups. Treating the swapped genes as truly differential features allows for benchmarking the sensitivity and specificity of the different DD workflows. As described in Section 4.4, these analyses where performed for three cell types (T4 naive cell, B memory cells and non-classical myeloid cells), on different sample sizes, and each analysis was performed five times on a different randomly selected subset of samples.

### 4.7. Performance measures

We assess the performance of the eight different DD workflows by visualizing the true positive rate (TPR) against the false discovery rate (FDR), which are computed according to the following definitions:

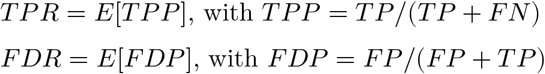

where FN, FP and TP denote the number of false negatives, false positives and true positives, respectively. The TPR is the expected value of the true positive proportion (TPP), which is a measure of sensitivity, while FDR is the expected value of the false discovery proportion (FDP), which is a measure of specificity. In Section 2.2, the expected value of the TPP and FDP is estimated by taking the average of the TPP and FDP over the five replicates that were generated for each simulated dataset, as described in Section 4.6, respectively.

### 4.8. Simulation for stage-wise analysis

In the simulations described above in Section 4.6, DD signal between groups is introduced by swapping counts between genes in one of the mock treatment groups. This, however, also introduces a DE signal. In this section, we introduce three other ad-hoc simulation strategies, that allow for simulating genes with different properties. Note that for each of these DD and/or DE signal simulation strategies, we avoid creating systematic differences between the mock treatment groups, such as differences in sequencing depth, by altering an equal number of genes in each mock treatment group.

The first strategy puts the expression values of the 25% of cells with the lowest non-zero expression values in one of the mock treatment groups to zero. This strategy will generate DD signal between the treatment groups, as the detection of the target gene is reduced in only one treatment group. While this strategy will also generate a DE signal, i.e. a shift in mean expression between the treatment groups, this signal will be relatively small, because only the lowest expression values are set to zero. We expect this signal to mimic real-life situations, in which genes that are truly lowly expressed in the cell are often not detected by the sequencing technology.

The second strategy is similar to the one discussed above. However, the counts of the cells that are set to zero are uniformly distributed over the cells of the same patient that still express the gene. The effect on the DD signal is identical, but this strategy enforces that the within-group average expression of the target gene is not altered. However, note that this strategy will result in increased variability in the expression data, and may result in artificially generating bimodal distributions for target genes.

In the third simulation strategy, we simply add a count of 1 to 25% of the cells in 1 treatment group that already expressed the gene. This way, we generate a DE signal without introducing DD.

Finally, note that for all simulation strategies we consider a setting with two mock treatment groups, and that changes for each gene are randomly applied either to the first or second mock treatment group, to avoid systematically altering the library size of cells in either treatment group.

### 4.9. COVID case study

We performed a DE analysis on the B-cells of the COVID case study by Stephenson et al. (2021). To ensure robustness, patient-cell type combinations with fewer than 20 cells were removed from the analysis. Specifically, B cells, class-switched memory B cells, immature B cells, naive B cells, and unswitched memory B cells were analyzed due to insufficient healthy patient data for IgA plasma cells, IgG plasma cells, IgM plasma cells and plasmablasts. Patients from Cambridge and Newcastle sequencing sites were retained, while patients from the Sanger sequencing site were excluded since the number of patients in each disease severity stratum was low and there were no healthy patients. Gene-level filtering was then applied for each cell type separately, removing genes with fewer than three UMIs in any given pseudobulk sample or fewer than five counts in total across donor pseudobulk samples. Healthy patients were analyzed alongside those with mild, moderate, critical or severe COVID. To account for possible confounders, sequencing site and gender were included as covariates. A DD workflow (edgeR_NB_optim) and a pseudobulk gene expression analysis with edgeR were applied to the data, and statistical significance was assessed at the 5% FDR level.

### 4.10. Lupus case study

The data used in this study is derived from a study on systemic lupus erythematosus (SLE) (Perez et al., 2022). We applied several filtering steps to address potential confounders. Specifically, we focused on European women, the largest demographic group, and selected samples from six specific batches (“dmx_YS-JY-20_pool3”, “dmx_YS-JY-20_pool4”, “dmx_YS-JY-21_pool1”, “dmx_YS-JY-21_pool2”, “dmx_YS-JY-22_pool5” and “dmx_YS-JY-22_pool6”). We retained data from three cell types (T4 naive cells, B memory cells and non-classical myeloid cells) as described in Section 4.4. Gene-level filtering was then applied for each cell type separately, retaining only genes with expression in at least 200 cells per cell type. Patient age and batch were included as covariates to account for potential confounders. A DD workflow (edgeR_NB_optim) and a pseudobulk DE analysis with edgeR were applied to the data, and statistical significance was assessed at the 5% FDR level.

### 4.11. Code availability

The source code to reproduce our analyses and figures is available from https://github.com/statOmics/DD_benchmarks for code related to our mock and performance benchmarks, and from https://github.com/statOmics/DD_cases for code related to our COVID and lupus case studies, respectively. These repositories contain all information to run our analyses from scratch. For snapshots of these repositories after running all the scripts, containing all input, intermediate data objects and output, we refer to our Zenodo repository https://doi.org/10.5281/zenodo.10391097 (Gilis et al.).

### 4.12. Data availability

The data that serves as input for our benchmarks and case studies, as well as intermediate data objects and final output data are available from Zenodo https://doi.org/10.5281/zenodo.10391097 (Gilis et al.).

This Zenodo repository project contains the following data objects:

- **Lupus_benchmark.zip:** folder containing the source code, intermediate objects and output for the mock and performance benchmarks of the eight different DD workflows on the lupus data. The folder structure is identical to that of the `benchmarks/lupus` folder of our companion github repository https://github.com/statOmics/DD_benchmarks.
- **Lupus-n_patients_benchmark.zip:** folder containing the source code, intermediate objects and output for the mock and performance benchmarks of the eight different DD workflows on the lupus data, where we test the effect of sample size on the type I error control and performance. The folder structure is identical to that of the `benchmarks/lupus-n_patients` folder of our companion github repository https://github.com/statOmics/DD_benchmarks.
- **Stagewise_benchmark.zip:** folder containing the source code, intermediate objects and output for the mock and performance benchmarks of the eight different DD workflows on the lupus data, where we test the stage-wise testing procedure. The folder structure is identical to that of the `benchmarks/lupus-n_patients` folder of our companion github repository https://github.com/statOmics/DD_benchmarks.
- **Covid_benchmark.zip:** folder containing the source code, intermediate objects and output for the mock and performance benchmarks of the eight different DD workflows on the COVID data. The folder structure is identical to that of the `benchmarks/covid` folder of our companion github repository https://github.com/statOmics/DD_benchmarks.
- **Renv_benchmarks.lock:** the metadata (e.g., version) about every R package that was used to perform the benchmark analyses.
- **Renv_benchmarks.zip:** a copy of the R libraries that were used to perform the benchmark analyses. Should be used combination with **Renv_benchmarks.lock** to reproduce the R environment that was used to perform the benchmark analyses.
- **Covid_case.zip:** folder containing the source code, intermediate objects and output for the COVID case study. The folder structure is identical to that of the `Covid` folder of our companion github repository https://github.com/statOmics/DD_cases.
- **Lupus_case.zip:** folder containing the source code, intermediate objects and output for the lupus case study. The folder structure is identical to that of the `Lupus` folder of our companion github repository https://github.com/statOmics/DD_cases.
- **Renv_cases.lock:** the metadata (e.g., version) about every R package that was used to perform the case study analyses.
- **Renv_cases.zip:** a copy of the R libraries that were used to perform the case study analyses. Should be used combination with **Renv_cases.lock** to reproduce the R environment that was used to perform the case study analyses.

All data are available under the terms of the Creative Commons Attribution 4.0 International license (CC-BY 4.0).

## 4.13. Acknowledgements

We would like to thankfully acknowledge Helena Crowell for a her insightful suggestions regarding Figure 3 of the manuscript. We also would like to acknowledge the students of the Statistical Genomics course, 2021/2022, Ghent University, who assisted us in assessing our initial implementation of the binomial GLM workflows for differential detection. Jeroen Gilis and Lieven Clement are supported by the Research Foundation Flanders (FWO), research grant No. G062219N, and Jeroen Gilis is further supported by FWO SB fellowship No. 3S037119. Milan Malfait has received research support from Johnson and Johnson Pharmaceuticals. Davide Risso was supported by the National Cancer Institute of the National Institutes of Health (U24CA180996) and by CZF2019-002443 from the Chan Zuckerberg Initiative DAF, an advised fund of Silicon Valley Community Foundation.

## Notes

https://doi.org/10.5281/zenodo.10391097

### Competing Interest Statement

The authors have declared no competing interest.

## References

Alfaro, E., Díaz-García, E., García-Tovar, S., Zamarrón, E., Mangas, A., Galera, R., López-Collazo, E., García-Rio, F., Cubillos-Zapata, C., 2022. Upregulated proteasome subunits in covid-19 patients: A link with hypoxemia, lymphopenia and inflammation. Biomolecules 12, 442. URL: https://www.mdpi.com/2218-273X/12/3/442, doi:10.3390/biom12030442

Arefinia, N., Yaghoubi, R., Ramezani, A., Farokhnia, M., Zadeh, A., Sarvari, J., 2023. Association of ifitm1 promoter methylation with severity of sars-cov-2 infection. Clinical Laboratory 69. URL: http://dx.doi.org/10.7754/Clin.Lab.2022.220622, doi:10.7754/clin.lab.2022.220622

Benjamini, Y., Hochberg, Y., 1995. Controlling the false discovery rate: A practical and powerful approach to multiple testing. Journal of the Royal Statistical Society: Series B (Methodological) 57, 289–300.

Van den Berge, K., Soneson, C., Robinson, M.D., Clement, L., 2017. stager: a general stage-wise method for controlling the gene-level false discovery rate in differential expression and differential transcript usage. Genome Biology 18, 151. URL: http://genomebiology.biomedcentral.com/articles/10.1186/s13059-017-1277-0, doi:10.1186/s13059-017-1277-0

Bergsneider, B., Bailey, E., Ahmed, Y., Gogineni, N., Huntley, D., Montano, X., 2021. Analysis of sars-cov-2 infection associated cell entry proteins ace2, cd147, ppia, and ppib in datasets from non sars-cov-2 infected neuroblastoma patients, as potential prognostic and infection biomarkers in neuroblastoma. Biochemistry and Biophysics Reports 27, 101081. URL: https://linkinghub.elsevier.com/retrieve/pii/S2405580821001758, doi:10.1016/j.bbrep.2021.101081

Bouland, G.A., Mahfouz, A., Reinders, M.J.T., 2021. Differential analysis of binarized single-cell rna sequencing data captures biological variation. NAR Genomics and Bioinformatics 3. URL: https://academic.oup.com/nargab/article/doi/10.1093/nargab/lqab118/6478878, doi:10.1093/nargab/lqab118

Chen, T.H., Chang, C.J., Hung, P.H., 2023. Possible pathogenesis and prevention of long covid: Sars-cov-2-induced mitochondrial disorder. International Journal of Molecular Sciences 24, 8034. URL: https://www.mdpi.com/1422-0067/24/9/8034, doi:10.3390/ijms24098034

Choudhary, S., Sreenivasulu, K., Mitra, P., Misra, S., Sharma, P., 2021. Role of genetic variants and gene expression in the susceptibility and severity of covid-19. Annals of Laboratory Medicine 41, 129–138. URL: http://annlabmed.org/journal/view.html?doi=10.3343/alm.2021.41.2.129, doi:10.3343/alm.2021.41.2.129

Crowell, H.L., Morillo Leonardo, S.X., Soneson, C., Robinson, M.D., 2022. Built on sand: the shaky foundations of simulating single-cell rna sequencing data. BioRxiv URL: http://biorxiv.org/lookup/doi/10.1101/2021.11.15.468676, doi:10.1101/2021.11.15.468676. DOI: 10.1101/2021.11.15.468676.

Crowell, H.L., Soneson, C., Germain, P.L., Calini, D., Collin, L., Raposo, C., Malhotra, D., Robinson, M.D., 2020. muscat detects subpopulation-specific state transitions from multi-sample multi-condition single-cell transcriptomics data. Nature Communications 11, 6077. URL: http://www.nature.com/articles/s41467-020-19894-4, doi:10.1038/s41467-020-19894-4

Finak, G., McDavid, A., Yajima, M., Deng, J., Gersuk, V., Shalek, A.K., Slichter, C.K., Miller, H.W., McElrath, M.J., Prlic, M., Linsley, P.S., Gottardo, R., 2015. Mast: a flexible statistical framework for assessing transcriptional changes and characterizing heterogeneity in single-cell rna sequencing data. Genome Biology 16, 278. URL: https://genomebiology.biomedcentral.com/articles/10.1186/s13059-015-0844-5, doi:10.1186/s13059-015-0844-5

Gilis, J., Perin, L., Malfait, M., Van den Berge, K., Takele Assefa, A., Verbist, B., Risso, D., Clement, L., . Datasets associated with the manuscript “differential detection workflows for multi-sample single-cell rna-seq data”. doi:10.5281/zenodo.10391097

Guarnieri, J.W., Dybas, J.M., Fazelinia, H., Kim, M.S., Frere, J., Zhang, Y., Albrecht, Y.S., Murdock, D.G., Angelin, A., Singh, L.N., Weiss, S.L., Best, S.M., Lott, M.T., Cope, H., Zaksas, V., Saravia-Butler, A., Meydan, C., Foox, J., Mozsary, C., Kidane, Y.H., Priebe, W., Emmett, M.R., Meller, R., Singh, U., Bram, Y., tenOever, B.R., Heise, M.T., Moorman, N.J., Madden, E.A., Taft-Benz, S.A., Anderson, E.J., Sanders, W.A., Dickmander, R.J., Baxter, V.K., Baylin, S.B., Wurtele, E.S., Moraes-Vieira, P.M., Taylor, D., Mason, C.E., Schisler, J.C., Schwartz, R.E., Beheshti, A., Wallace, D.C., 2022. Targeted Down Regulation Of Core Mitochondrial Genes During SARS-CoV-2 Infection. Technical Report. URL: http://biorxiv.org/lookup/doi/10.1101/2022.02.19.481089, doi:10.1101/2022.02.19.481089. DOI: 10.1101/2022.02.19.481089

Henriques-Pons, A., Beghini, D.G., Silva, V.D.S., Iwao Horita, S., Silva, F., 2022. Pulmonary mesenchymal stem cells in mild cases of covid-19 are dedicated to proliferation; in severe cases, they control inflammation, make cell dispersion, and tissue regeneration. Frontiers in Immunology 12, 780900. URL: https://www.frontiersin.org/articles/10.3389/fimmu.2021.780900/full, doi:10.3389/fimmu.2021.780900

Hicks, S.C., Townes, F.W., Teng, M., Irizarry, R.A., 2018. Missing data and technical variability in single-cell rna-sequencing experiments. Biostatistics 19, 562–578. URL: https://academic.oup.com/biostatistics/article/19/4/562/4599254, doi:10.1093/biostatistics/kxx053

Huang, R., Chen, W., Zhao, X., Ma, Y., Zhou, Q., Chen, J., Zhang, M., Zhao, D., Hou, Y., He, C., Wu, Y., 2023. Genome-wide characterization of alternative splicing in blood cells of covid-19 and respiratory infections of relevance. Virologica Sinica 38, 309–312. URL: https://linkinghub.elsevier.com/retrieve/pii/S1995820X2300007X, doi:10.1016/j.virs.2023.01.007

Jeyananthan, P., 2022. Comprehensive machine learning analysis on the phenotypes of covid-19 patients using transcriptome data. Arab Gulf Journal of Scientific Research, 79–137URL: https://agjsr.agu.edu.bh/publications/paper/1326, doi:10.51758/AGJSR-S2-2021-0023

Khatoon, F., Prasad, K., Kumar, V., 2020. Neurological manifestations of covid-19: available evidences and a new paradigm. Journal of NeuroVirology 26, 619–630. URL: https://link.springer.com/10.1007/s13365-020-00895-4, doi:10.1007/s13365-020-00895-4

Korthauer, K.D., Chu, L.F., Newton, M.A., Li, Y., Thomson, J., Stewart, R., Kendziorski, C., 2016. A statistical approach for identifying differential distributions in single-cell rna-seq experiments. Genome Biology 17, 222. URL: https://genomebiology.biomedcentral.com/articles/10.1186/s13059-016-1077-y, doi:10.1186/s13059-016-1077-y

Lata, S., Mishra, R., Arya, R.P., Arora, P., Lahon, A., Banerjea, A.C., Sood, V., 2022. Where all the roads meet? a crossover perspective on host factors regulating sars-cov-2 infection. Journal of Molecular Biology 434, 167403. URL: https://linkinghub.elsevier.com/retrieve/pii/S0022283621006409, doi:10.1016/j.jmb.2021.167403

Li, C.x., Chen, J., Lv, S.k., Li, J.h., Li, L.l., Hu, X., 2021. Whole-transcriptome rna sequencing reveals significant differentially expressed mrnas, mirnas, and lncrnas and related regulating biological pathways in the peripheral blood of covid-19 patients. Mediators of Inflammation 2021, 1–22. URL: https://www.hindawi.com/journals/mi/2021/6635925/, doi:10.1155/2021/6635925

Love, M.I., Huber, W., Anders, S., 2014. Moderated estimation of fold change and dispersion for rna-seq data with deseq2. Genome Biology 15, 550. URL: http://genomebiology.biomedcentral.com/articles/10.1186/s13059-014-0550-8, doi:10.1186/s13059-014-0550-8

Lun, A.T.L., Marioni, J.C., 2017. Overcoming confounding plate effects in differential expression analyses of single-cell rna-seq data. Biostatistics 18, 451–464. URL: https://academic.oup.com/biostatistics/article/18/3/451/2970368, doi:10.1093/biostatistics/kxw055

Lund, S.P., Nettleton, D., McCarthy, D.J., Smyth, G.K., 2012. Detecting differential expression in rna-sequence data using quasi-likelihood with shrunken dispersion estimates. Statistical Applications in Genetics and Molecular Biology 11. URL: https://www.degruyter.com/document/doi/10.1515/1544-6115.1826/html, doi:10.1515/1544-6115.1826

Mellett, L., Khader, S.A., 2022. S100a8/a9 in covid-19 pathogenesis: Impact on clinical outcomes. Cytokine and Growth Factor Reviews 63, 90–97. URL: https://linkinghub.elsevier.com/retrieve/pii/S1359610121000769, doi:10.1016/j.cytogfr.2021.10.004

Melnichuk, N., Liashko, V., Kashuba, V., Tkachuk, Z., 2022. Candidates-biomarkers for the express test-system to diagnosis of sars-cov-2-induced immunopathology URL: https://febs.onlinelibrary.wiley.com/doi/epdf/10.1002/2211-5463.13440.

Momeni, M., Rashidifar, M., Balam, F.H., Roointan, A., Gholaminejad, A., 2023. A comprehensive analysis of gene expression profiling data in covid-19 patients for discovery of specific and differential blood biomarker signatures. Scientific Reports 13, 5599. URL: https://www.nature.com/articles/s41598-023-32268-2, doi:10.1038/s41598-023-32268-2

Mukund, K., Nayak, P., Ashokkumar, C., Rao, S., Almeda, J., Betancourt-Garcia, M.M., Sindhi, R., Subramaniam, S., 2021. Immune response in severe and non-severe coronavirus disease 2019 (covid-19) infection: A mechanistic landscape. Frontiers in Immunology 12, 738073. URL: https://www.frontiersin.org/articles/10.3389/fimmu.2021.738073/full, doi:10.3389/fimmu.2021.738073

Murphy, A.E., Skene, N.G., 2022. A balanced measure shows superior performance of pseudobulk methods in single-cell rna-sequencing analysis. Nature Communications 13, 7851. URL: https://www.nature.com/articles/s41467-022-35519-4, doi:10.1038/s41467-022-35519-4

Park, J.H., Lee, H.K., 2020. Re-analysis of single cell transcriptome reveals that the nr3c1-cxcl8-neutrophil axis determines the severity of covid-19. Frontiers in Immunology 11, 2145. URL: https://www.frontiersin.org/article/10.3389/fimmu.2020.02145/full, doi:10.3389/fimmu.2020.02145

Perez, R.K., Gordon, M.G., Subramaniam, M., Kim, M.C., Hartoularos, G.C., Targ, S., Sun, Y., Ogorodnikov, A., Bueno, R., Lu, A., Thompson, M., Rappoport, N., Dahl, A., Lanata, C.M., Matloubian, M., Maliskova, L., Kwek, S.S., Li, T., Slyper, M., Waldman, J., Dionne, D., Rozenblatt-Rosen, O., Fong, L., Dall”Era, M., Balliu, B., Regev, A., Yazdany, J., Criswell, L.A., Zaitlen, N., Ye, C.J., 2022. Single-cell rna-seq reveals cell type–specific molecular and genetic associations to lupus. Science 376, eabf1970. URL: https://www.science.org/doi/10.1126/science.abf1970, doi:10.1126/science.abf1970

Qi, F., Zhang, W., Huang, J., Fu, L., Zhao, J., 2021. Single-cell rna sequencing analysis of the immunometabolic rewiring and immunopathogenesis of coronavirus disease 2019. Frontiers in Immunology 12, 651656. URL: https://www.frontiersin.org/articles/10.3389/fimmu.2021.651656/full, doi:10.3389/fimmu.2021.651656

Qiu, P., 2020. Embracing the dropouts in single-cell rna-seq analysis. Nature Communications 11, 1169. URL: https://www.nature.com/articles/s41467-020-14976-9, doi:10.1038/s41467-020-14976-9

Rashmi, P., Sur, S., Sigdel, T.K., Boada, P., Schroeder, A.W., Damm, I., Kretzler, M., Hodgin, J., Hartoularos, G., Jimmie Ye, C., Sarwal, M.M., 2022. Multiplexed droplet single-cell sequencing (mux-seq) of normal and transplant kidney. American Journal of Transplantation 22, 876–885. URL: https://linkinghub.elsevier.com/retrieve/pii/S1600613522081400, doi:10.1111/ajt.16871

Robinson, M.D., McCarthy, D.J., Smyth, G.K., 2010. edger a bioconductor package for differential expression analysis of digital gene expression data. Bioinformatics 26, 139–140. URL: https://academic.oup.com/bioinformatics/article-lookup/doi/10.1093/bioinformatics/btp616, doi:10.1093/bioinformatics/btp616

Smyth, G.K., 2004. Linear models and empirical bayes methods for assessing differential expression in microarray experiments. Statistical Applications in Genetics and Molecular Biology 3, 1–25. URL: https://www.degruyter.com/document/doi/10.2202/1544-6115.1027/html, doi:10.2202/1544-6115.1027

Soneson, C., Robinson, M.D., 2018. Bias, robustness and scalability in single-cell differential expression analysis. Nature Methods 15, 255–261. URL: http://www.nature.com/articles/nmeth.4612, doi:10.1038/nmeth.4612

Squair, J.W., Gautier, M., Kathe, C., Anderson, M.A., James, N.D., Hutson, T.H., Hudelle, R., Qaiser, T., Matson, K.J.E., Barraud, Q., Levine, A.J., La Manno, G., Skinnider, M.A., Courtine, G., 2021. Confronting false discoveries in singlecell differential expression. Nature Communications 12, 5692. URL: https://www.nature.com/articles/s41467-021-25960-2, doi:10.1038/s41467-021-25960-2

Stephenson, E., Reynolds, G., Botting, R.A., Calero-Nieto, F.J., Morgan, M.D., Tuong, Z.K., Bach, K., Sungnak, W., Worlock, K.B., Yoshida, M., Kumasaka, N., Kania, K., Engelbert, J., Olabi, B., Spegarova, J.S., Wilson, N.K., Mende, N., Jardine, L., Gardner, L.C.S., Goh, I., Horsfall, D., McGrath, J., Webb, S., Mather, M.W., Lindeboom, R.G.H., Dann, E., Huang, N., Polanski, K., Prigmore, E., Gothe, F., Scott, J., Payne, R.P., Baker, K.F., Hanrath, A.T., Schim Van Der Loeff, I.C.D., Barr, A.S., Sanchez-Gonzalez, A., Bergamaschi, L., Mescia, F., Barnes, J.L., Kilich, E., De Wilton, A., Saigal, A., Saleh, A., Janes, S.M., Smith, C.M., Gopee, N., Wilson, C., Coupland, P., Coxhead, J.M., Kiselev, V.Y., Van Dongen, S., Bacardit, J., King, H.W., Rostron, A.J., Simpson, A.J., Hambleton, S., Laurenti, E., Lyons, P.A., Meyer, K.B., Nikolić, M.Z., Duncan, C.J.A., Smith, K.G.C., Teichmann, S.A., Clatworthy, M.R., Marioni, J.C., Göttgens, B., Haniffa, M., 2021. Single-cell multi-omics analysis of the immune response in covid-19. Nature Medicine 27, 904–916. URL: https://www.nature.com/articles/s41591-021-01329-2, doi:10.1038/s41591-021-01329-2

Tiberi, S., Crowell, H.L., Samartsidis, P., Weber, L.M., Robinson, M.D., 2023. distinct: a novel approach to differential distribution analyses. The Annals of Applied Statistics doi:10.1101/2020.11.24.394213

Townes, F.W., Hicks, S.C., Aryee, M.J., Irizarry, R.A., 2019. Feature selection and dimension reduction for single-cell rna-seq based on a multinomial model. Genome Biology 20, 295. URL: https://genomebiology.biomedcentral.com/articles/10.1186/s13059-019-1861-6, doi:10.1186/s13059-019-1861-6

Wen, W., Su, W., Tang, H., Le, W., Zhang, X., Zheng, Y., Liu, X., Xie, L., Li, J., Ye, J., Dong, L., Cui, X., Miao, Y., Wang, D., Dong, J., Xiao, C., Chen, W., Wang, H., 2020. Immune cell profiling of covid-19 patients in the recovery stageby single-cell sequencing. Cell Discovery 6, 31. URL: https://www.nature.com/articles/s41421-020-0168-9, doi:10.1038/s41421-020-0168-9

Wilson, D.J., 2019. The harmonic mean p-value for combining dependent tests. Proceedings of the National Academy of Sciences 116, 1195–1200. URL: https://pnas.org/doi/full/10.1073/pnas.1814092116, doi:10.1073/pnas.1814092116

Yang, X., Chi, H., Wu, M., Wang, Z., Lang, Q., Han, Q., Wang, X., Liu, X., Li, Y., Wang, X., Huang, N., Bi, J., Liang, H., Gao, Y., Zhao, Y., Feng, N., Yang, S., Wang, T., Xia, X., Ge, L., 2022. Discovery and characterization of sars-cov-2 reactive and neutralizing antibodies from humanized camousehg mice through rapid hybridoma screening and high-throughput single-cell v(d)j sequencing. Frontiers in Immunology 13, 992787. URL: https://www.frontiersin.org/articles/10.3389/fimmu.2022.992787/full, doi:10.3389/fimmu.2022.992787

Yuka, S.A., Yilmaz, A., 2021. Effect of sars-cov-2 infection on host competing endogenous rna and mirna network. PeerJ 9, e12370. URL: https://peerj.com/articles/12370, doi:10.7717/peerj.12370

Zhang, M., Liu, S., Miao, Z., Han, F., Gottardo, R., Sun, W., 2022. Ideas: individual level differential expression analysis for single-cell rna-seq data. Genome Biology 23, 33. URL: https://genomebiology.biomedcentral.com/articles/10.1186/s13059-022-02605-1, doi:10.1186/s13059-022-02605-1

Zhao, X.N., You, Y., Cui, X.M., Gao, H.X., Wang, G.L., Zhang, S.B., Yao, L., Duan, L.J., Zhu, K.L., Wang, Y.L., Li, L., Lu, J.H., Wang, H.B., Fan, J.F., Zheng, H.W., Dai, E.H., Tian, L.Y., Ma, M.J., 2021. Single-cell immune profiling reveals distinct immune response in asymptomatic covid-19 patients. Signal Transduction and Targeted Therapy 6, 342. URL: https://www.nature.com/articles/s41392-021-00753-7, doi:10.1038/s41392-021-00753-7

Zimmerman, K.D., Espeland, M.A., Langefeld, C.D., 2021. A practical solution to pseudoreplication bias in single-cell studies. Nature Communications 12, 738. URL: https://www.nature.com/articles/s41467-021-21038-1, doi:10.1038/s41467-021-21038-1

