## Supplementary Figures and Tables for "Differential detection workflows for multi-sample single-cell RNA-seq data"

### Supplementary figures for the manuscript; “Differential detection workflows for multi-patient single-cell RNA-seq data”

Jeroen Gilis[a,b,c,d], Laura Perin[a,e], Milan Malfait[b], Koen Van den Berge[f], Alemu Takele Assefa[f], Bie Verbist[f], Davide Risso[e,g], Lieven Clement[b,c,*]

[a] *These authors contributed equally,*

[b] *Applied Mathematics, Computer science and Statistics, Ghent University, Ghent, 9000, Belgium,*

[c] *Bioinformatics Institute, Ghent University, Ghent, 9000, Belgium,*

[d] *Data Mining and Modeling for Biomedicine, VIB Flemish Institute for Biotechnology, Ghent, 9000, Belgium,*

[e] *Department of Statistical Sciences, University of Padova, Padova, Italy,*

[f] *Statistics and Decision Sciences, Johnson and Johnson Innovative Medicine, Beerse, Belgium,*

[g] *Padua Center for Network Medicine, University of Padova, Padova, Italy,*

Supplementary table 1: **Top 10 differentially detected genes that are not differentially expressed**. The first column displays the top 10 DD genes, unique to Naive B cells in moderately diseased patients. The second column showcases the top 10 DD genes, specific to Naive B cells in critically diseased patients. The third column highlights the top 10 DD genes exclusive to unswitched B memory cells in moderately diseased patients. Finally, the fourth column exhibits the top 10 DD genes identified in unswitched B memory cells in critically diseased patients.

| Naive Moderate | | Naive critical | | Unswitched Moderate | | Unswitched critical | |
|---|---|---|---|---|---|---|---|
| ATP5F1A | 4.37e-07 | IFITM1 | 0.0002 | PSME2 | 5.38e-06 | EIF5A | 1.51e-07 |
| IGHV6-1 | 3.23e-05 | IGHV6-1 | 0.0004 | IGKC | 9.38e-06 | DDX21 | 1.08e-06 |
| ACTG1 | 4.35e-05 | MCM5 | 0.0005 | PMAIP1 | 2.42e-05 | IGKC | 1.32e-06 |
| GDI2 | 4.63e-05 | SNX6 | 0.0006 | KIAA0040 | 3.36e-05 | COX6A1 | 1.36e-05 |
| HSPA8 | 0.0001 | MYC | 0.0008 | NFKBID | 3.42e-05 | CD27 | 3.24e-05 |
| HMGB2 | 0.0002 | S100A8 | 0.001 | DDX21 | 5.15e-05 | RHOA | 3.48e-05 |
| LITAF | 0.0002 | IGVK1D-8 | 0.001 | CD1C | 7.32e-05 | CD1C | 3.57e-05 |
| IGLV1-51 | 0.0003 | | | ZNF581 | 7.42e-05 | ZNF581 | 3.80e-05 |
| SHMT2 | 0.0003 | | | TNIP2 | 0.0001 | TNIP2 | 4.43e-05 |
| PKM | 0.0005 | | | PSME2 | 0.0001 | PSME2 | 4.45e-05 |

*Corresponding author

Supplementary table 2: **Overlap Between the individual DE and DD Analysis Results with the stage2 DD and DE Results. Column 1:** an identifier for the comparison (contrast) for easy referencing. **Column 2:** the cell type (B: B cell, CsMB: Class-switched memory B cell, ImmB: immature B cell, NaiveB: naive B cell and UMB: unswitched memory B cell). **Column 3:** the COVID status of the samples that are compared to the healthy samples. **Column 4:** the number of differentially expressed genes identified by the edgeR analysis at the 5% false discovery rate (FDR) level. **Column 5:** the number of differentially expressed genes identified in the second stage of the stage-wise testing procedure. **Column 6:** the overlap between columns 4 and 5. **Column 7:** the number of differentially detected genes identified by edgeR_NB_optim at the 5% FDR level. **Column 8:** the number of differentially expressed genes identified in the second stage of the stage-wise testing procedure. **Column 9:** the overlap between columns 8 and 9.

| Compa -rison | Cell type | COVID status | DE | DE stage2 | Overlap DE - DE stage2 | DD | DD stage2 | Overlap DD - DD stage2 |
|---|---|---|---|---|---|---|---|---|
| 1 | B | Asymptomatic | 0 | 0 | 0 | 0 | 0 | 0 |
| 2 | B | Mild | 0 | 0 | 0 | 2 | 1 | 1 |
| 3 | B | Moderate | 7 | 5 | 5 | 3 | 3 | 3 |
| 4 | B | Severe | 1 | 1 | 1 | 1 | 1 | 1 |
| 5 | B | Critical | 0 | 0 | 0 | 6 | 0 | 0 |
| 6 | CsMB | Asymptomatic | 0 | 0 | 0 | 0 | 0 | 0 |
| 7 | CsMB | Mild | 84 | 81 | 79 | 8 | 33 | 8 |
| 8 | CsMB | Moderate | 339 | 326 | 322 | 191 | 213 | 191 |
| 9 | CsMB | Severe | 138 | 138 | 133 | 50 | 72 | 50 |
| 10 | CsMB | Critical | 4 | 6 | 4 | 11 | 10 | 10 |
| 11 | ImmB | Asymptomatic | 1 | 1 | 1 | 1 | 1 | 1 |
| 12 | ImmB | Mild | 48 | 55 | 46 | 36 | 40 | 36 |
| 13 | ImmB | Moderate | 329 | 297 | 297 | 110 | 154 | 110 |
| 14 | ImmB | Severe | 72 | 74 | 71 | 43 | 52 | 43 |
| 15 | ImmB | Critical | 0 | 9 | 0 | 27 | 19 | 19 |
| 16 | NaiveB | Asymptomatic | 0 | 0 | 0 | 0 | 0 | 0 |
| 17 | NaiveB | Mild | 251 | 229 | 228 | 135 | 161 | 135 |
| 18 | NaiveB | Moderate | 2437 | 2239 | 2239 | 1417 | 1580 | 1409 |
| 19 | NaiveB | Severe | 1676 | 1474 | 1474 | 558 | 812 | 555 |
| 20 | NaiveB | Critical | 1307 | 1125 | 1125 | 224 | 453 | 224 |
| 21 | UMB | Asymptomatic | 0 | 0 | 0 | 0 | 0 | 0 |
| 22 | UMB | Mild | 14 | 21 | 14 | 19 | 21 | 19 |
| 23 | UMB | Moderate | 119 | 159 | 119 | 164 | 162 | 154 |
| 24 | UMB | Severe | 78 | 84 | 78 | 11 | 30 | 11 |
| 25 | UMB | Critical | 0 | 3 | 0 | 50 | 14 | 14 |

Supplementary table 3: **GSEA Results for Unswitched B Memory Cells of Critically Ill Patients. Column 1:** The gene ontology (GO) terms of the processes that are significantly enriched. **Column 2:** Ranking of the GO terms when performing a GSEA on the significant DD genes of the stage-wise analysis. **Column 3:** P-values for the GO terms when performing a GSEA on the significant DD genes of the stage-wise analysis. Note that for this comparison, no genes were found to be differentially expressed. Hence, the GSEA was not performed for the DE and stage-wise analysis results.

| GO Term | Rank DD stage2 | pvalue DD stage2 |
|---|---|---|
| Oxydative phosphorylation | 1 | 8.0e-04 |
| Regulation of cell shape | 2 | 1.0e-03 |
| Positive regulation of translational termination | 3 | 1.5e-03 |
| Proton transmembrane transport | 4 | 1.8e-03 |
| Aerobic respiration | 5 | 1.9e-03 |
| Cortical microtubule organization | 6 | 2.2e-03 |
| Intestinal D-glucose absorption | 7 | 2.2e-03 |

| GO Term | Rank DD stage2 | pvalue DD stage2 |
|---|---|---|
| Intestinal hexose absorption | 8 | 2.2e-03 |
| Membrane to membrane docking | 9 | 2.2e-03 |
| Positive regulation of translational elongation | 10 | 2.2e-03 |

Supplementary table 4: **GSEA Results for Naive B Cells of Moderately Ill Patients. Column 1:** The gene ontology (GO) terms of the processes that are significantly enriched. **Column 2:** Ranking of the GO terms when performing a GSEA on the significant DD genes of the stage-wise analysis. **Column 3:** Ranking of the GO terms when performing a GSEA on the significant DE genes of the stage-wise analysis. **Column 4:** Ranking of the GO terms when performing a GSEA on the significant genes from the omnibus test of the stage-wise analysis. **Column 5:** P-values for the GO terms when performing a GSEA on the significant DD genes of the stage-wise analysis. **Column 6:** P-values for the GO terms when performing a GSEA on the significant DE genes of the stage-wise analysis. **Column 7:** P-values for the GO terms when performing a GSEA on the significant genes from the omnibus test of the stage-wise analysis.

| GO Term | Rank DD stage2 | Rank DE stage2 | Rank Omnibus | pvalue DD stage2 | pvalue DE stage2 | pvalue Omnibus |
|---|---|---|---|---|---|---|
| Biological process | 1 | 10 | 9 | 2.4e-19 | 4.2e-33 | 3.0e-34 |
| Cellular process | 2 | 9 | 8 | 1.7e-18 | 8.3e-34 | 1.6e-35 |
| Cellular macromolecule metabolic process | 3 | 3 | 3 | 1.7e-14 | 4.1e-44 | 1.2e-43 |
| Mitochondrial translation | 4 | 23 | 23 | 1.7e-12 | 3.7e-20 | 3.2e-21 |
| Cellular metabolic process | 5 | 12 | 11 | 2.5e-12 | 1.2e-29 | 1.4e-31 |
| Metabolic process | 6 | 13 | 13 | 1.2e-11 | 1.7e-29 | 3.4e-31 |
| Macromolecule metabolic process | 7 | 18 | 17 | 2.5e-10 | 7.5e-27 | 4.5e-28 |
| Nitrogen compound metabolic process | 8 | 14 | 14 | 2.9e-10 | 4.0e-29 | 1.1e-30 |
| Mitochondrial gene expression | 9 | 26 | 26 | 4.9e-10 | 4.7e-19 | 7.2e-20 |
| Protein metabolic process | 10 | 15 | 16 | 1.3e-09 | 4.4e-28 | 1.2e-28 |
| Translation | 11 | 1 | 1 | 1.5e-09 | 1.4e-52 | 4.0e-52 |
| Peptide biosynthetic process | 12 | 2 | 2 | 2.8e-09 | 8.8e-51 | 2.9e-50 |
| Amide biosynthetic process | 27 | 4 | 4 | 1.6e-07 | 2.5e-43 | 1.4e-42 |
| Peptide metabolic process | 31 | 5 | 5 | 3.5e-07 | 9.7e-43 | 6.9e-42 |
| Cytoplasmic translation | 623 | 6 | 6 | 3.3e-02 | 1.0e-39 | 1.6e-39 |
| Cellular macromolecule biosynthetic process | 35 | 7 | 7 | 1.1e-06 | 1.5e-37 | 2.8e-36 |
| Cellular amide metabolic process | 63 | 8 | 10 | 1.4e-05 | 5.3e-34 | 8.1e-33 |

Supplementary table 5: **GSEA Results for Naive B Cells of Critically Ill Patients. Column 1:** The gene ontology (GO) terms of the processes that are significantly enriched. **Column 2:** Ranking of the GO terms when performing a GSEA on the significant DD genes of the stage-wise analysis. **Column 3:** Ranking of the GO terms when performing a GSEA on the significant DE genes of the stage-wise analysis. **Column 4:** Ranking of the GO terms when performing a GSEA on the significant genes from the omnibus test of the stage-wise analysis. **Column 5:** P-values for the GO terms when performing a GSEA on the significant DD genes of the stage-wise analysis. **Column 6:** P-values for the GO terms when performing a GSEA on the significant DE genes of the stage-wise analysis. **Column 7:** P-values for the GO terms when performing a GSEA on the significant genes from the omnibus test of the stage-wise analysis.

| GO Term | Rank DD stage2 | Rank DE stage2 | Rank Omnibus | pvalue DD stage2 | pvalue DE stage2 | pvalue Omnibus |
|---|---|---|---|---|---|---|
| regulation of cell cycle process | 1 | 168 | 159 | 1.2e-08 | 1.5e-04 | 1.0e-04 |
| mitotic cell cycle phase transition | 2 | 107 | 100 | 1.4e-08 | 6.5e-06 | 3.3e-06 |
| chromosome organization | 3 | 96 | 59 | 2.2e-08 | 2.9e-06 | 2.4e-07 |
| mitotic cell cycle | 4 | 110 | 109 | 3.5e-08 | 8.9e-06 | 6.5e-06 |

| GO Term | Rank DD stage2 | Rank DE stage2 | Rank Omni-bus | pvalue DD stage2 | pvalue DE stage2 | pvalue Omni-bus |
|---|---|---|---|---|---|---|
| mitotic cell cycle process | 5 | 100 | 96 | 6.2e-08 | 3.8e-06 | 2.6e-06 |
| cell cycle | 6 | 184 | 173 | 6.4e-08 | 2.4e-04 | 1.4e-04 |
| cell cycle phase transition | 7 | 150 | 135 | 1.5e-07 | 7.8e-05 | 4.6e-05 |
| kinetochore organization | 8 | 105 | 110 | 1.6e-07 | 6.4e-06 | 7.1e-06 |
| regulation of chromosome segregation | 9 | 116 | 117 | 2.6e-07 | 1.0e-05 | 1.3e-05 |
| mitotic nuclear division | 10 | 62 | 66 | 3.5e-07 | 3.6e-07 | 5.0e-07 |
| cellular macromolecule metabolic process | 22 | 1 | 1 | 2.4e-06 | 1.5e-33 | 7.2e-34 |
| translation | 297 | 2 | 2 | 1.8e-02 | 8.0e-33 | 2.7e-32 |
| peptide biosynthetic process | 253 | 3 | 3 | 1.2e-02 | 3.0e-32 | 1.0e-31 |
| peptide metabolic process | 144 | 4 | 4 | 2.8e-03 | 4.0e-31 | 1.5e-30 |
| organonitrogen compound biosynthetic process | 345 | 5 | 5 | 2.3e-02 | 9.1e-31 | 5.3e-30 |
| cellular macromolecule biosynthetic process | 809 | 6 | 6 | 7.8e-02 | 2.3e-28 | 9.2e-28 |
| amide biosynthetic process | 542 | 7 | 7 | 4.7e-02 | 7.8e-28 | 2.6e-27 |
| cellular amide metabolic process | 291 | 8 | 8 | 1.7e-02 | 3.0e-25 | 1.1e-24 |
| organonitrogen compound metabolic process | 121 | 9 | 12 | 1.5e-03 | 1.6e-22 | 1.6e-21 |
| cellular process | 14 | 10 | 9 | 7.9e-07 | 5.1e-22 | 3.7e-22 |
| cellular nitrogen compound metabolic process | 45 | 12 | 10 | 7.4e-05 | 4.4e-21 | 1.2e-21 |

Supplementary table 6: **Comparison of Individual DD and DE analysis GSEA results and stage2 GSEA Results for Unswitched Memory B Cells of Moderately Ill Patients. Column 1:** The gene ontology (GO) terms of the processes that are significantly enriched. **Column 2:** Ranking of the GO terms when performing a GSEA on the significant genes of the DD analysis. **Column 3:** Ranking of the GO terms when performing a GSEA on the significant DD genes of the stage-wise analysis. **Column 4:** P-values for the GO terms when performing a GSEA on the significant genes of the DD analysis. **Column 5:** P-values for the GO terms when performing a GSEA on the significant DD genes of the stage-wise analysis.**Column 6:** Ranking of the GO terms when performing a GSEA on the significant genes of the DE analysis. **Column 7:** Ranking of the GO terms when performing a GSEA on the significant DE genes of the stage-wise analysis. **Column 8:** P-values for the GO terms when performing a GSEA on the significant genes of the DE analysis. **Column 9:** P-values for the GO terms when performing a GSEA on the significant DE genes of the stage-wise analysis.

| GO Term | Rank DD | Rank DD stage2 | pvalue DD | pvalue DD stage2 | Rank DE | Rank DE stage2 | pvalue DE | pvalue DE stage2 |
|---|---|---|---|---|---|---|---|---|
| Response to virus | 1 | 1 | 2.6e-12 | 4.1e-13 | 12 | 19 | 3.1e-08 | 1.3e-08 |
| Defense response to symbiont | 2 | 3 | 1.4e-10 | 1.9e-11 | 21 | 23 | 3.7e-06 | 6.0e-07 |
| Defense response to virus | 3 | 4 | 1.4e-10 | 1.9e-11 | 22 | 24 | 3.7e-06 | 6.0e-07 |
| Immune response | 4 | 2 | 3.2e-10 | 8.0e-12 | 19 | 13 | 3.9e-07 | 6.1e-10 |
| Response to interferon-alpha | 5 | 9 | 6.0e-10 | 8.1e-10 | 34 | 32 | 8.2e-05 | 5.1e-06 |
| Response to stress | 6 | 5 | 1.2e-09 | 1.1e-10 | 47 | 56 | 3.5e-04 | 1.4e-04 |
| Immune system process | 7 | 6 | 2.9e-09 | 1.6e-10 | 27 | 21 | 1.8e-05 | 1.3e-07 |
| Response to other organism | 8 | 7 | 3.0e-08 | 5.4e-10 | 29 | 27 | 2.8e-05 | 2.7e-06 |
| Response to external biotic stimulus | 9 | 8 | 3.2e-08 | 5.7e-10 | 30 | 28 | 2.9e-05 | 2.9e-06 |
| Viral genome replication | 10 | 16 | 4.8e-08 | 7.1e-08 | 33 | 26 | 5.6e-05 | 1.6e-06 |
| Response to biotic stimulus | 11 | 10 | 5.5e-08 | 1.1e-09 | 32 | 30 | 4.2e-05 | 4.6e-06 |
| Cytoplasmic translation | 1164 | 1281 | 2.3e-01 | 2.5e-01 | 1 | 1 | 3.9e-21 | 3.7e-29 |
| Translation | 658 | 748 | 8.7e-02 | 1.0e-01 | 2 | 2 | 2.8e-17 | 1.3e-23 |
| Peptide biosynthetic process | 704 | 824 | 9.9e-02 | 1.2e-01 | 3 | 3 | 6.3e-17 | 4.0e-23 |
| Amide biosynthetic process | 983 | 1131 | 1.7e-01 | 2.0e-01 | 4 | 4 | 2.2e-15 | 5.9e-21 |
| Peptide metabolic process | 1093 | 1246 | 2.1e-01 | 2.4e-01 | 5 | 5 | 7.8e-15 | 3.5e-20 |

| GO Term | Rank DD | Rank DD stage2 | pvalue DD | pvalue DD stage2 | Rank DE | Rank DE stage2 | pvalue DE | pvalue DE stage2 |
|---|---|---|---|---|---|---|---|---|
| Cellular macromolecule biosynthetic process | 1235 | 1404 | 2.6e-01 | 3.0e-01 | 6 | 6 | 2.2e-12 | 1.1e-17 |
| Cellular amide metabolic process | 1576 | 1717 | 4.2e-01 | 4.7e-01 | 7 | 7 | 2.4e-12 | 1.1e-16 |
| Gene expression | 34 | 51 | 9.7e-05 | 3.4e-04 | 8 | 9 | 9.2e-12 | 3.2e-14 |
| Cellular nitrogen compound biosynthetic process | 51 | 54 | 3.4e-04 | 3.9e-04 | 9 | 10 | 1.2e-10 | 1.5e-12 |
| Macromolecule biosynthetic process | 88 | 96 | 1.9e-03 | 2.1e-03 | 10 | 11 | 1.4e-10 | 1.8e-12 |
| Organonitrogen compound biosynthetic process | 1453 | 1345 | 3.6e-01 | 2.8e-01 | 11 | 8 | 9.4e-10 | 2.4e-14 |

Supplementary table 7: **Comparison of Individual DD and DE analysis GSEA results and stage2 GSEA Results for Unswitched Memory B Cells of Critically Ill Patients. Column 1:** The gene ontology (GO) terms of the processes that are significantly enriched. **Column 2:** Ranking of the GO terms when performing a GSEA on the significant genes of the DD analysis. **Column 3:** Ranking of the GO terms when performing a GSEA on the significant DD genes of the stage-wise analysis. **Column 4:** P-values for the GO terms when performing a GSEA on the significant genes of the DD analysis. **Column 5:** P-values for the GO terms when performing a GSEA on the significant DD genes of the stage-wise analysis.

| GO Term | Rank DD | Rank DD stage2 | pvalue DD | pvalue DD stage2 |
|---|---|---|---|---|
| Oxidative phosphorylation | 1 | 1 | 2.0e-07 | 8.0e-04 |
| Aerobic respiration | 2 | 5 | 1.9e-06 | 1.9e-03 |
| Cellular respiration | 3 | 15 | 6.9e-06 | 3.3e-03 |
| Response to interferon-beta | 4 | 10440 | 7.4e-06 | 1 |
| Proton motive force-driven mitochondrial ATP synthesis | 5 | 59 | 1.2e-05 | 1.5e-02 |
| Proton motive force-driven ATP synthesis | 6 | 75 | 3.1e-05 | 2.0e-02 |
| Energy derivation by oxidation of organic compounds | 7 | 33 | 3.5e-05 | 6.2e-03 |
| Proton transmembrane transport | 8 | 4 | 5.1e-05 | 1.8e-03 |
| Filopodium assembly | 9 | 88 | 6.4e-05 | 2.5e-02 |
| Localization within membrane | 10 | 87 | 1.0e-04 | 2.4e-02 |
| Regulation of cell shape | 90 | 2 | 1.0e-02 | 1.0e-03 |
| Positive regulation of translational termination | 59 | 3 | 4.7e-03 | 1.5e-03 |
| Cortical microtubule organization | 68 | 6 | 7.0e-03 | 2.2e-03 |
| Intestinal D-glucose absorption | 69 | 7 | 7.0e-03 | 2.2e-03 |
| Intestinal hexose absorption | 70 | 8 | 7.0e-03 | 2.2e-03 |
| Membrane to membrane docking | 71 | 9 | 7.0e-03 | 2.2e-03 |
| Positive regulation of translational elongation | 73 | 10 | 7.0e-03 | 2.2e-03 |

Supplementary table 8: **Comparison of Individual DD and DE analysis GSEA results and stage2 GSEA Results for Naive B Cells of Moderately Ill Patients. Column 1:** The gene ontology (GO) terms of the processes that are significantly enriched. **Column 2:** Ranking of the GO terms when performing a GSEA on the significant genes of the DD analysis. **Column 3:** Ranking of the GO terms when performing a GSEA on the significant DD genes of the stage-wise analysis. **Column 4:** P-values for the GO terms when performing a GSEA on the significant genes of the DD analysis. **Column 5:** P-values for the GO terms when performing a GSEA on the significant DD genes of the stage-wise analysis.**Column 6:** Ranking of the GO terms when performing a GSEA on the significant genes of the DE analysis. **Column 7:** Ranking of the GO terms when performing a GSEA on the significant DE genes of the stage-wise analysis. **Column 8:** P-values for the GO terms when performing a GSEA on the significant genes of the DE analysis. **Column 9:** P-values for the GO terms when performing a GSEA on the significant DE genes of the stage-wise analysis.

| GO Term | Rank DD | Rank DD stage2 | pvalue DD | pvalue DD stage2 | Rank DE | Rank DE stage2 | pvalue DE | pvalue DE stage2 |
|---|---|---|---|---|---|---|---|---|
| Cellular process | 1 | 2 | 1.2e-17 | 1.7e-18 | 8 | 9 | 2.8e-34 | 8.3e-34 |
| Biological process | 2 | 1 | 3.8e-17 | 2.4e-19 | 9 | 10 | 1.4e-33 | 4.2e-33 |
| Cellular macromolecule metabolic process | 3 | 3 | 8.2e-15 | 1.7e-14 | 3 | 3 | 9.1e-43 | 4.1e-44 |
| Cellular metabolic process | 4 | 5 | 1.3e-12 | 2.5e-12 | 13 | 12 | 7.1e-30 | 1.2e-29 |
| Mitochondrial translation | 5 | 4 | 5.6e-12 | 1.7e-12 | 22 | 23 | 9.0e-20 | 3.7e-20 |
| Metabolic process | 6 | 6 | 4.3e-11 | 1.2e-11 | 12 | 13 | 4.7e-30 | 1.7e-29 |
| Macromolecule metabolic process | 7 | 7 | 4.8e-10 | 2.5e-10 | 17 | 18 | 3.3e-27 | 7.5e-27 |
| Mitochondrial gene expression | 8 | 9 | 1.1e-09 | 4.9e-10 | 26 | 26 | 1.7e-18 | 4.7e-19 |
| Translation | 9 | 11 | 1.3e-09 | 1.5e-09 | 1 | 1 | 2.1e-50 | 1.4e-52 |
| Nitrogen compound metabolic process | 10 | 8 | 1.3e-09 | 2.9e-10 | 14 | 14 | 7.2e-30 | 4.0e-29 |
| Protein metabolic process | 12 | 10 | 1.3e-08 | 1.3e-09 | 16 | 15 | 1.9e-27 | 4.4e-28 |
| Peptide biosynthetic process | 11 | 12 | 2.1e-09 | 2.8e-09 | 2 | 2 | 1.4e-48 | 8.8e-51 |
| Amide biosynthetic process | 18 | 27 | 8.5e-08 | 1.6e-07 | 4 | 4 | 1.4e-41 | 2.5e-43 |
| Peptide metabolic process | 22 | 31 | 1.6e-07 | 3.5e-07 | 5 | 5 | 1.8e-41 | 9.7e-43 |
| Cytoplasmic translation | 428 | 623 | 1.5e-02 | 3.3e-02 | 6 | 6 | 1.1e-38 | 1.0e-39 |
| Cellular macromolecule biosynthetic process | 24 | 35 | 2.3e-07 | 1.1e-06 | 7 | 7 | 1.4e-35 | 1.5e-37 |
| Cellular amide metabolic process | 51 | 63 | 7.4e-06 | 1.4e-05 | 10 | 8 | 3.2e-33 | 5.3e-34 |

Supplementary table 9: **Comparison of Individual DD and DE analysis GSEA results and stage2 GSEA Results for Naive B Cells of Critically Ill Patients. Column 1:** The gene ontology (GO) terms of the processes that are significantly enriched. **Column 2:** Ranking of the GO terms when performing a GSEA on the significant genes of the DD analysis. **Column 3:** Ranking of the GO terms when performing a GSEA on the significant DD genes of the stage-wise analysis. **Column 4:** P-values for the GO terms when performing a GSEA on the significant genes of the DD analysis. **Column 5:** P-values for the GO terms when performing a GSEA on the significant DD genes of the stage-wise analysis.**Column 6:** Ranking of the GO terms when performing a GSEA on the significant genes of the DE analysis. **Column 7:** Ranking of the GO terms when performing a GSEA on the significant DE genes of the stage-wise analysis. **Column 8:** P-values for the GO terms when performing a GSEA on the significant genes of the DE analysis. **Column 9:** P-values for the GO terms when performing a GSEA on the significant DE genes of the stage-wise analysis.

| GO Term | Rank DD | Rank DD stage2 | pvalue DD | pvalue DD stage2 | Rank DE | Rank DE stage2 | pvalue DE | pvalue DE stage2 |
|---|---|---|---|---|---|---|---|---|
| Chromosome organization | 1 | 3 | 8.4e-07 | 2.2e-08 | 83 | 96 | 7.6e-07 | 2.9e-06 |
| Mitotic cell cycle | 2 | 4 | 1.6e-06 | 3.5e-08 | 121 | 110 | 1.3e-05 | 8.9e-06 |
| Regulation of cell cycle process | 3 | 1 | 2.0e-06 | 1.2e-08 | 213 | 168 | 2.9e-04 | 1.5e-04 |
| Chromosome condensation | 4 | 18 | 2.2e-06 | 1.7e-06 | 260 | 198 | 7.0e-04 | 3.0e-04 |
| Mitotic cell cycle process | 5 | 5 | 3.1e-06 | 6.2e-08 | 102 | 100 | 3.4e-06 | 3.8e-06 |

| GO Term | Rank DD | Rank DD stage2 | pvalue DD | pvalue DD stage2 | Rank DE | Rank DE stage2 | pvalue DE | pvalue DE stage2 |
|---|---|---|---|---|---|---|---|---|
| Mitotic chromosome condensation | 6 | 21 | 8.0e-06 | 2.3e-06 | 230 | 175 | 4.2e-04 | 2.0e-04 |
| Sulfur compound metabolic process | 7 | 89 | 8.6e-06 | 5.8e-04 | 297 | 249 | 1.4e-03 | 8.3e-04 |
| Mitotic cell cycle phase transition | 8 | 2 | 2.2e-05 | 1.4e-08 | 120 | 107 | 1.1e-05 | 6.5e-06 |
| Regulation of chromosome condensation | 9 | 64 | 2.8e-05 | 1.6e-04 | 376 | 326 | 3.7e-03 | 2.3e-03 |
| Cell cycle process | 10 | 20 | 3.2e-05 | 2.1e-06 | 234 | 225 | 4.8e-04 | 5.4e-04 |
| Cell cycle | 12 | 6 | 5.3e-05 | 6.4e-08 | 202 | 184 | 2.3e-04 | 2.4e-04 |
| Cell cycle phase transition | 11 | 7 | 4.4e-05 | 1.5e-07 | 147 | 150 | 3.9e-05 | 7.8e-05 |
| Kinetochore organization | 67 | 8 | 2.0e-03 | 1.6e-07 | 131 | 105 | 2.0e-05 | 6.4e-06 |
| Regulation of chromosome segregation | 15 | 9 | 8.0e-05 | 2.6e-07 | 86 | 116 | 1.0e-06 | 1.0e-05 |
| Mitotic nuclear division | 18 | 10 | 1.8e-04 | 3.5e-07 | 75 | 62 | 3.6e-07 | 3.6e-07 |
| Translation | 1911 | 297 | 3.9e-01 | 1.8e-02 | 1 | 2 | 9.8e-38 | 8.0e-33 |
| Peptide biosynthetic process | 1523 | 253 | 2.7e-01 | 1.2e-02 | 2 | 3 | 6.3e-37 | 3.0e-32 |
| Cellular macromolecule metabolic process | 37 | 22 | 8.0e-04 | 2.4e-06 | 3 | 1 | 7.6e-37 | 1.5e-33 |
| Peptide metabolic process | 917 | 144 | 1.2e-01 | 2.8e-03 | 4 | 4 | 3.8e-35 | 4.0e-31 |
| Organonitrogen compound biosynthetic process | 970 | 345 | 1.3e-01 | 2.3e-02 | 5 | 5 | 2.5e-33 | 9.1e-31 |
| Cellular macromolecule biosynthetic process | 2117 | 809 | 4.9e-01 | 7.8e-02 | 6 | 6 | 5.8e-32 | 2.3e-28 |
| Amide biosynthetic process | 1983 | 542 | 4.2e-01 | 4.7e-02 | 7 | 7 | 1.7e-31 | 7.8e-28 |
| Cellular amide metabolic process | 1089 | 291 | 1.5e-01 | 1.7e-02 | 8 | 8 | 3.2e-28 | 3.0e-25 |
| Cellular process | 28 | 14 | 5.4e-04 | 7.9e-07 | 9 | 10 | 9.3e-28 | 5.1e-22 |
| Cellular nitrogen compound metabolic process | 920 | 45 | 1.2e-01 | 7.4e-05 | 10 | 12 | 1.0e-26 | 4.4e-21 |
| Organonitrogen compound metabolic process | 259 | 121 | 2.4e-02 | 1.5e-03 | 14 | 9 | 2.0e-25 | 1.6e-22 |

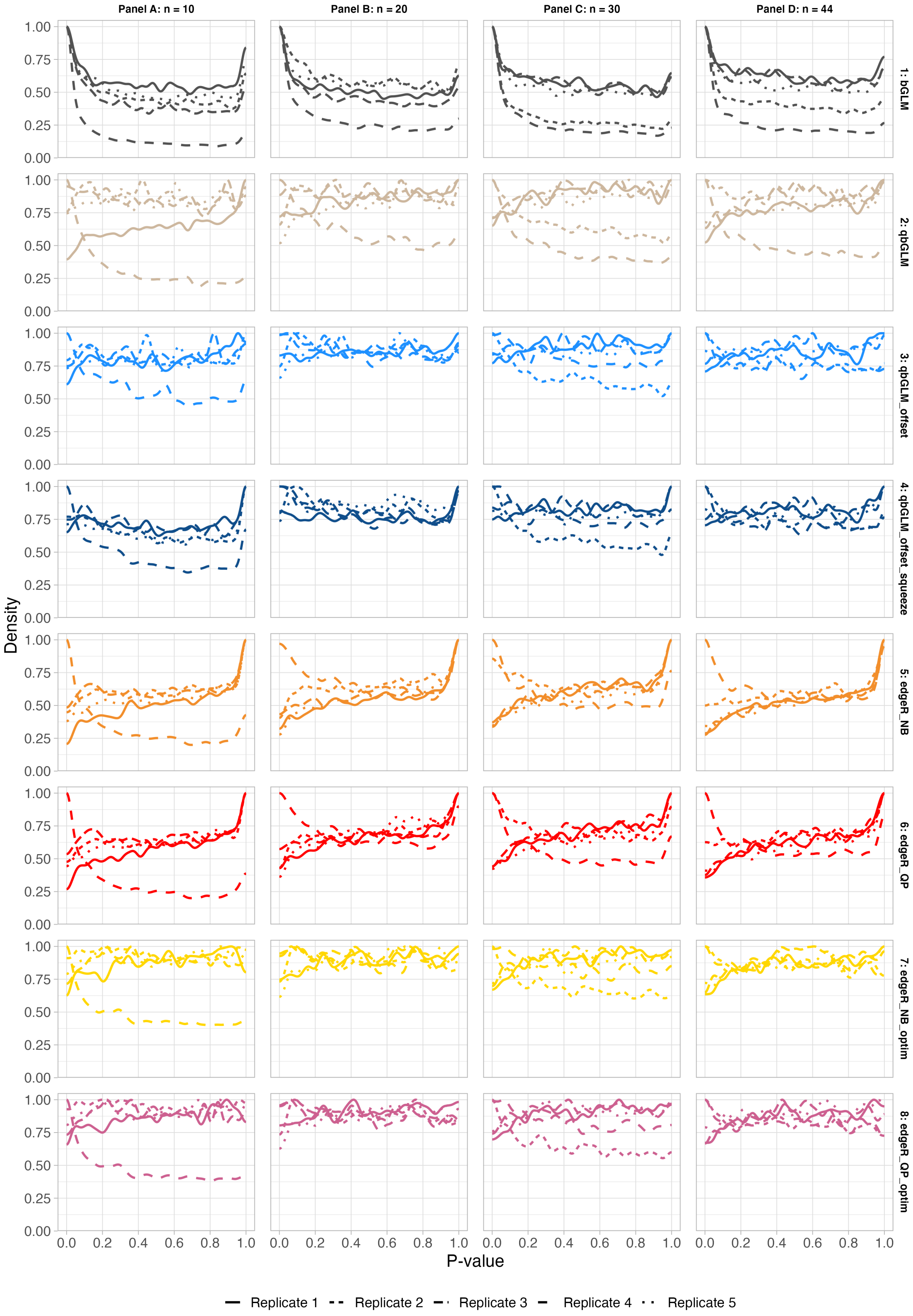


Supplementary figure 1: **Nominal p-value densities obtained from five mock simulation replicates based on the non-classical myeloid cells from the lupus dataset, pseudobulk data, stratified by method.** The sample size of each mock simulation is indicated in the column header, the name of the differential detection method is indicated in the rows.

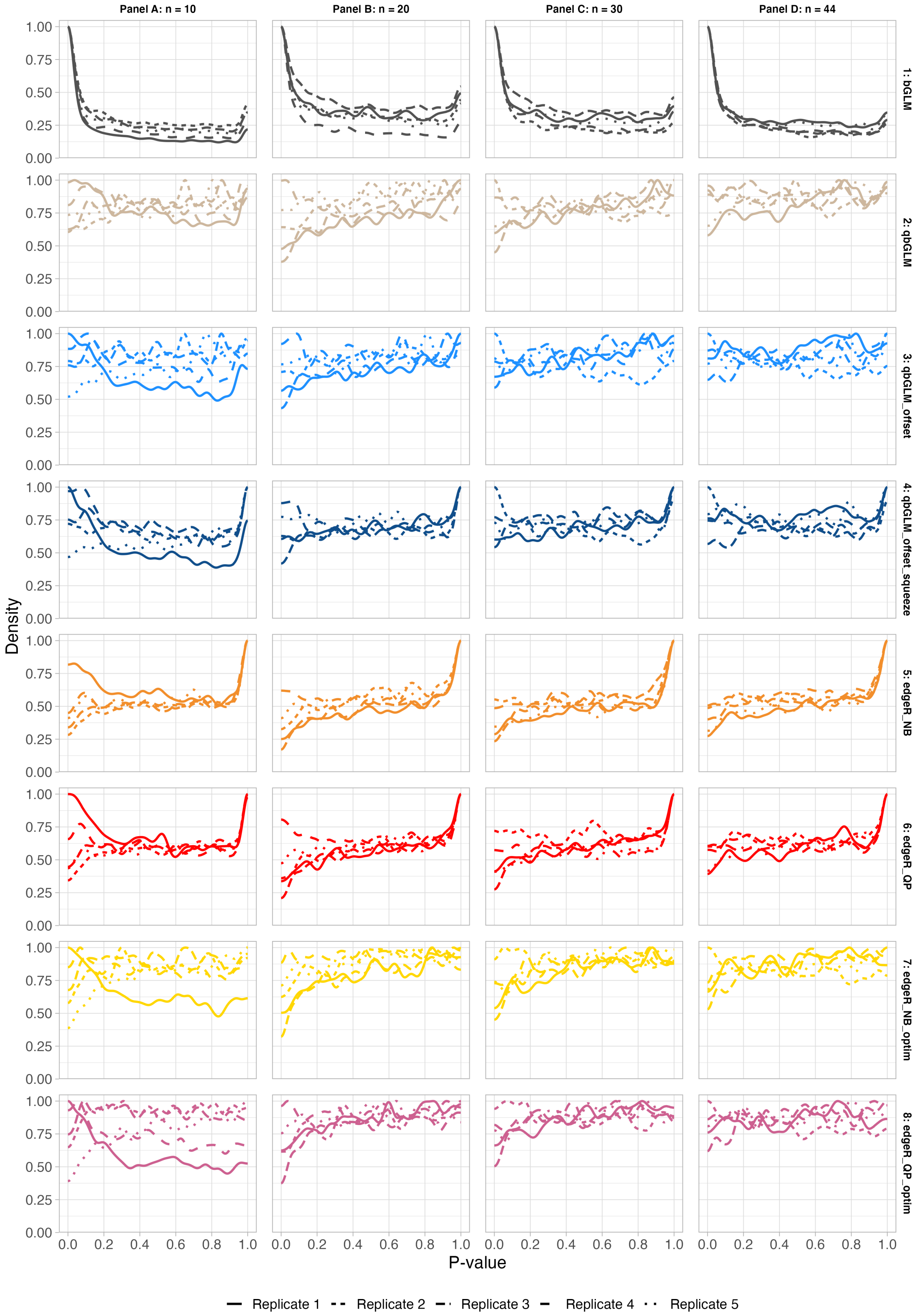


Supplementary figure 2: **Nominal p-value densities obtained from five mock simulation replicates based on the T4 naive cells from the lupus dataset, pseudobulk data, stratified by method.** The sample size of each mock simulation is indicated in the column header, the name of the differential detection method is indicated in the rows.

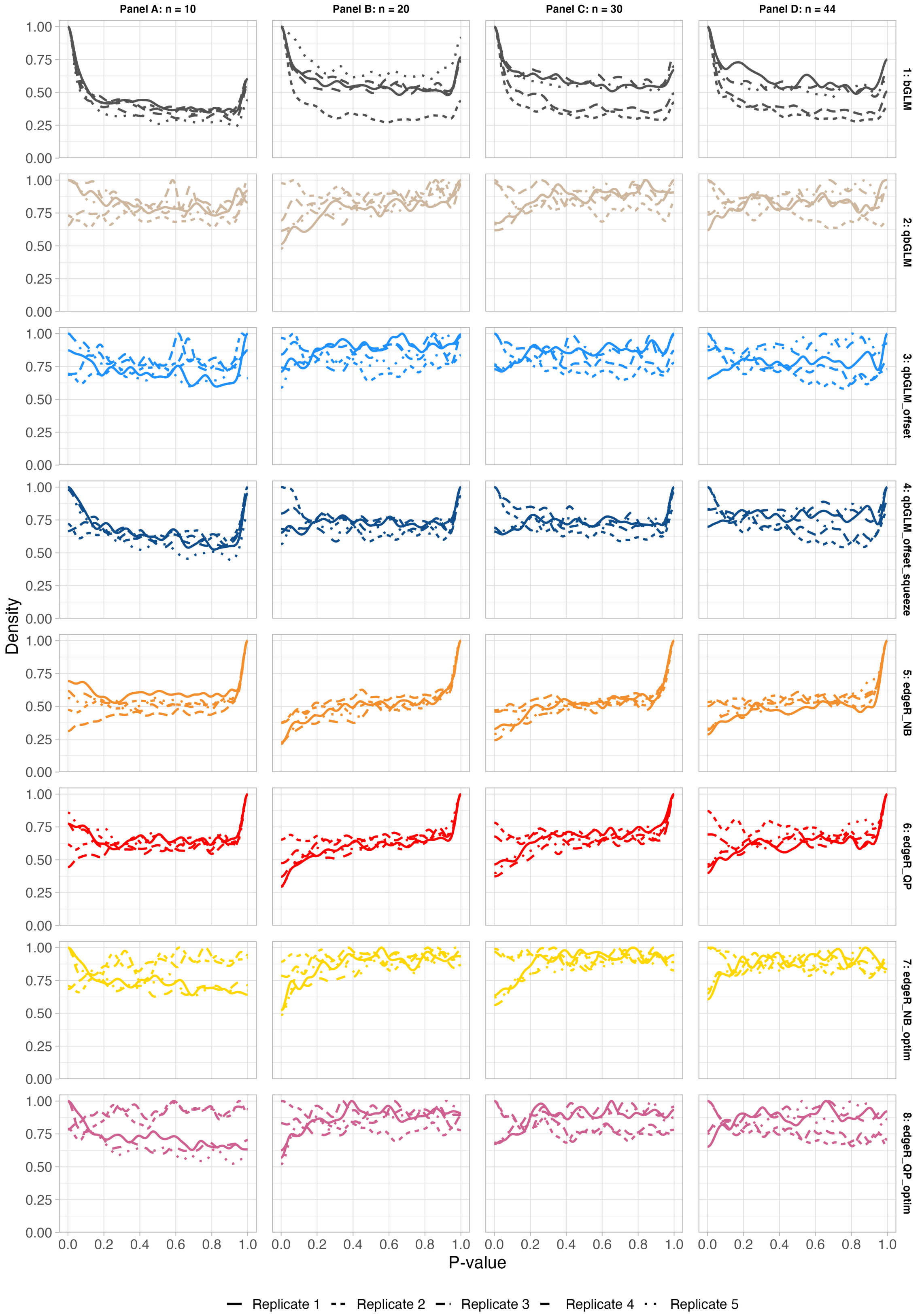


Supplementary figure 3: **Nominal p-value densities obtained from five mock simulation replicates based on the memory B cells from the lupus dataset, pseudobulk data, stratified by method.** The sample size of each mock simulation is indicated in the column header, the name of the differential detection method is indicated in the rows.

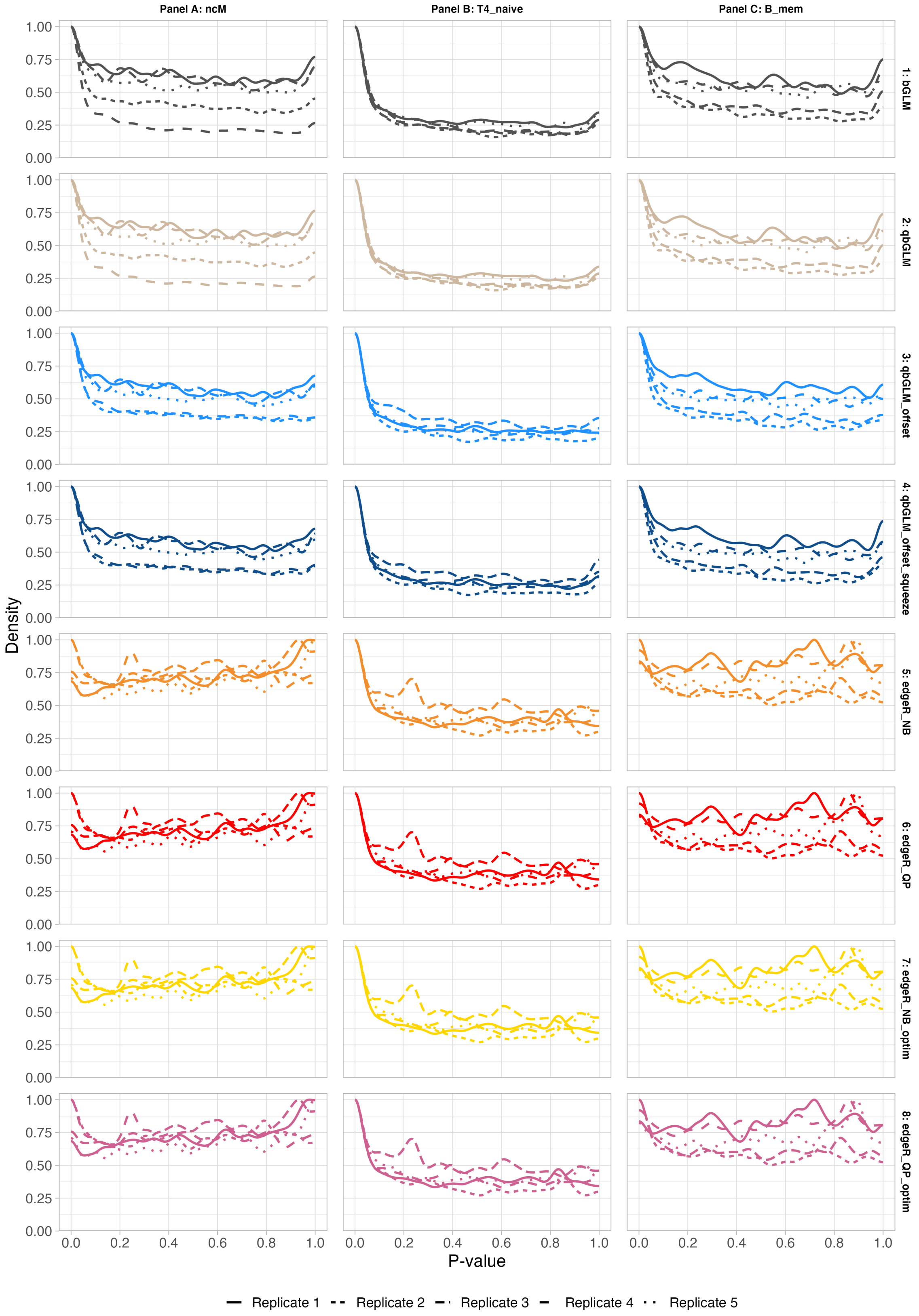


Supplementary figure 4: **Nominal p-value densities obtained from five mock simulation replicates based on the memory B cells from the lupus dataset, single-cell data, stratified by method.** These are the results on the three different cell types, as indicated in the columns, for the mock simulation with 22 versus 22 patients. For most cell types and methods, results are overly liberal. Note that the results are most liberal for the T4 naive cell type, which is the cell type with the largest number of cells per patient.

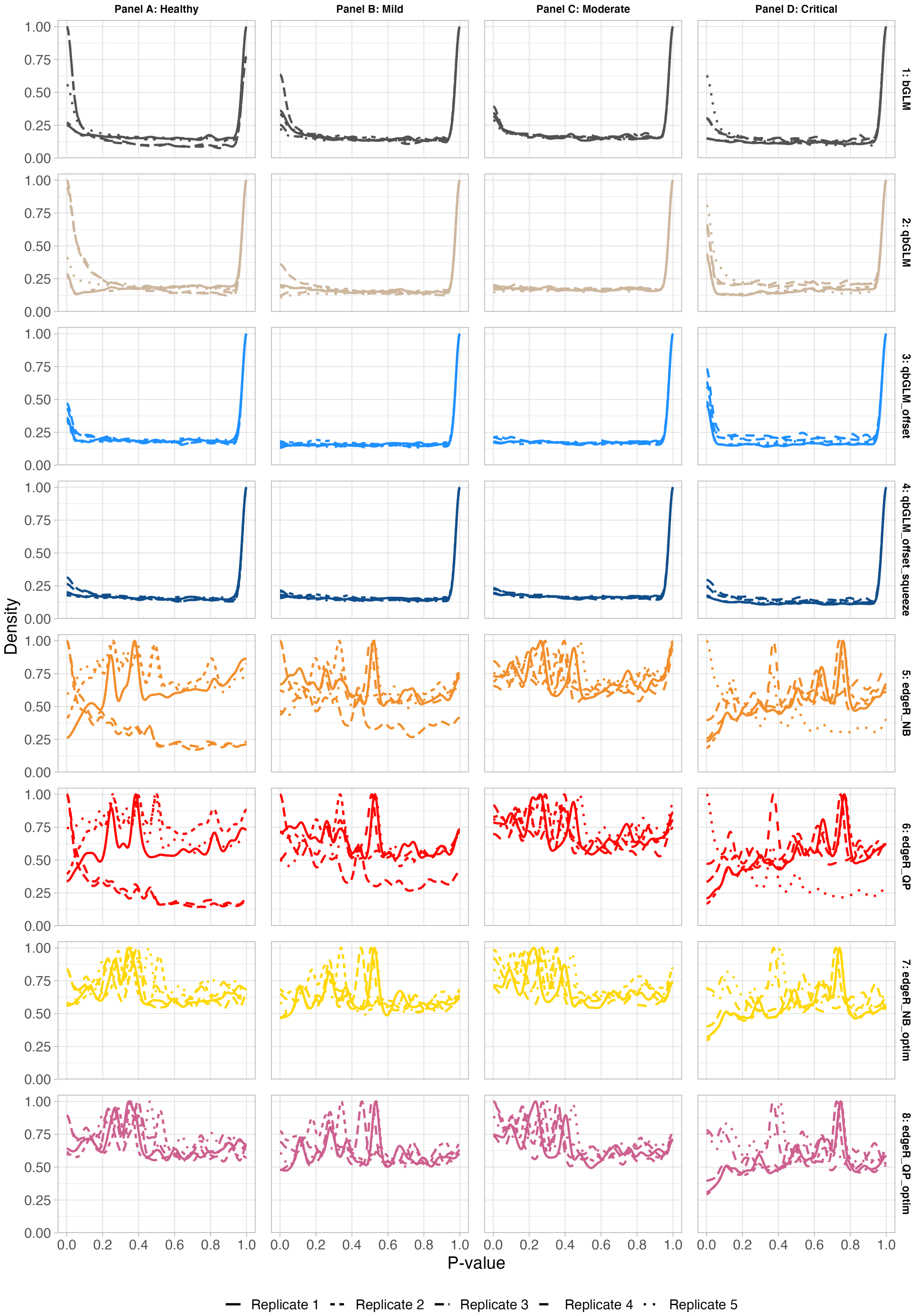


Supplementary figure 5: **Nominal p-value densities obtained from five mock simulation replicates based on the class switched memory B cells from the covid dataset, pseudobulk data, stratified by method.** The disease status of the patients on which the mock simulation is based is indicated in the column header, the name of the differential detection method is indicated in the rows.

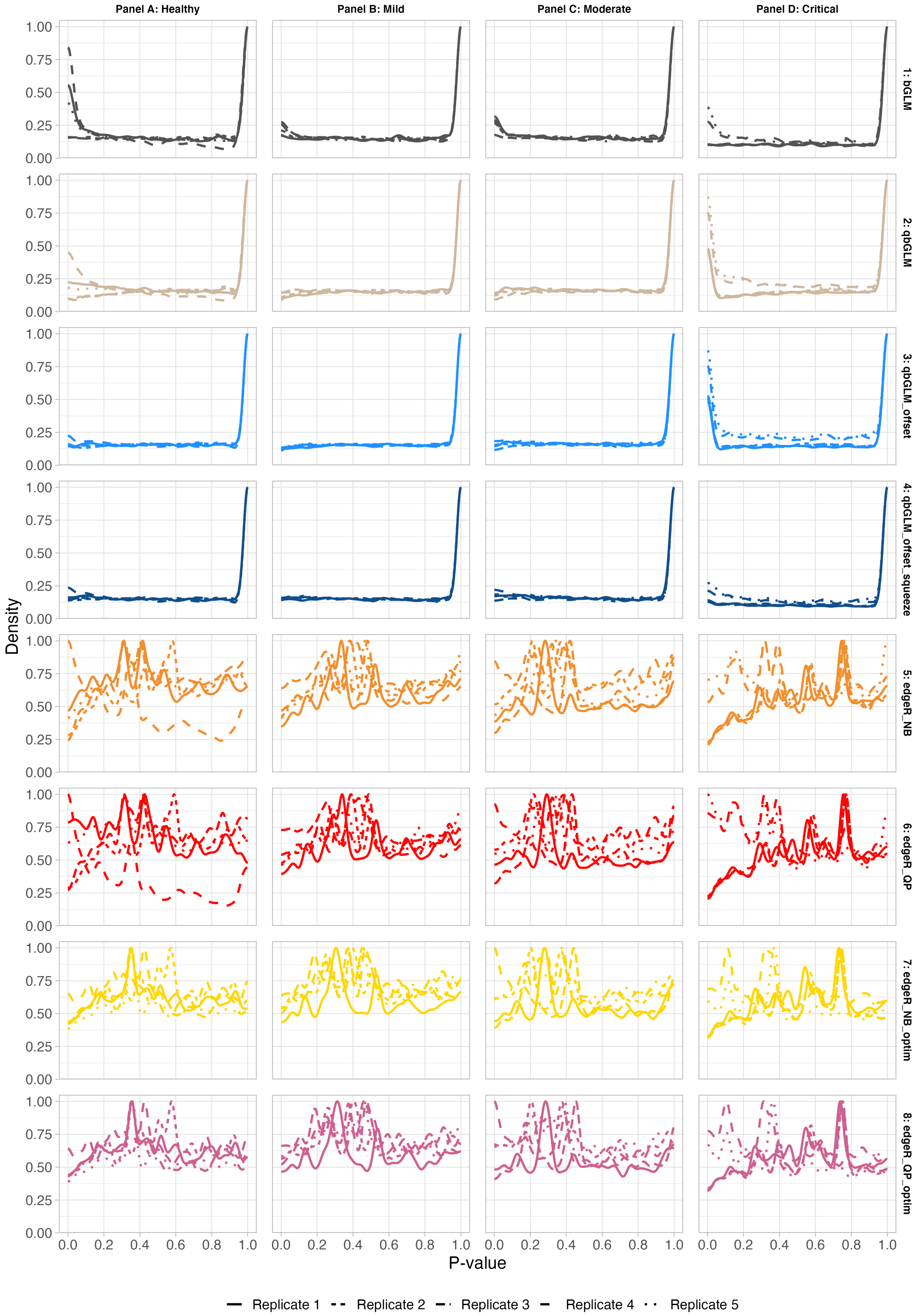


Supplementary figure 6: **Nominal p-value densities obtained from five mock simulation replicates based on the immature B cells from the covid dataset, pseudobulk data, stratified by method.** The disease status of the patients on which the mock simulation is based is indicated in the column header, the name of the differential detection method is indicated in the rows.

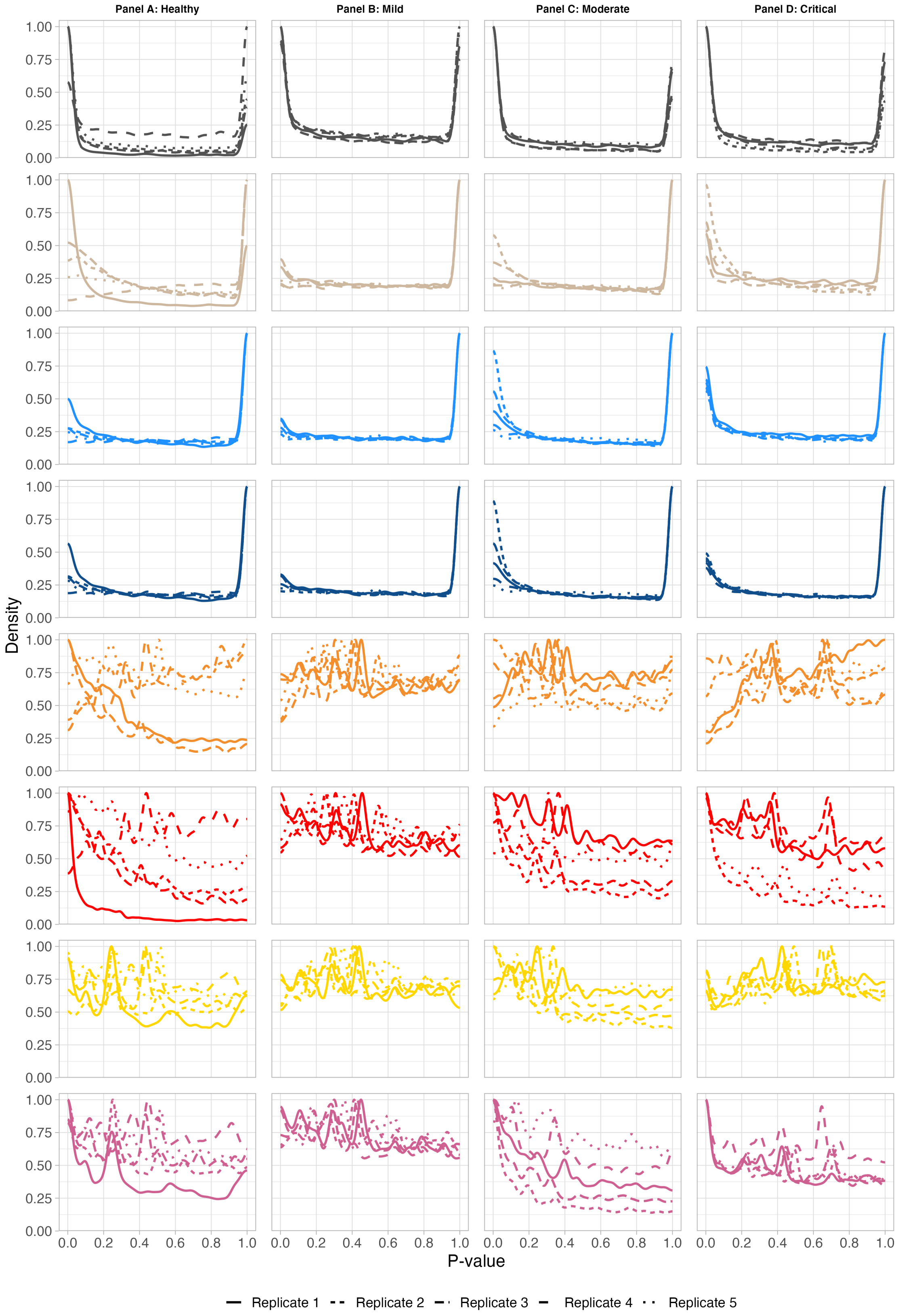


Supplementary figure 7: **Nominal p-value densities obtained from five mock simulation replicates based on the naive B cells from the covid dataset, pseudobulk data, stratified by method.** The disease status of the patients on which the mock simulation is based is indicated in the column header, the name of the differential detection method is indicated in the rows.

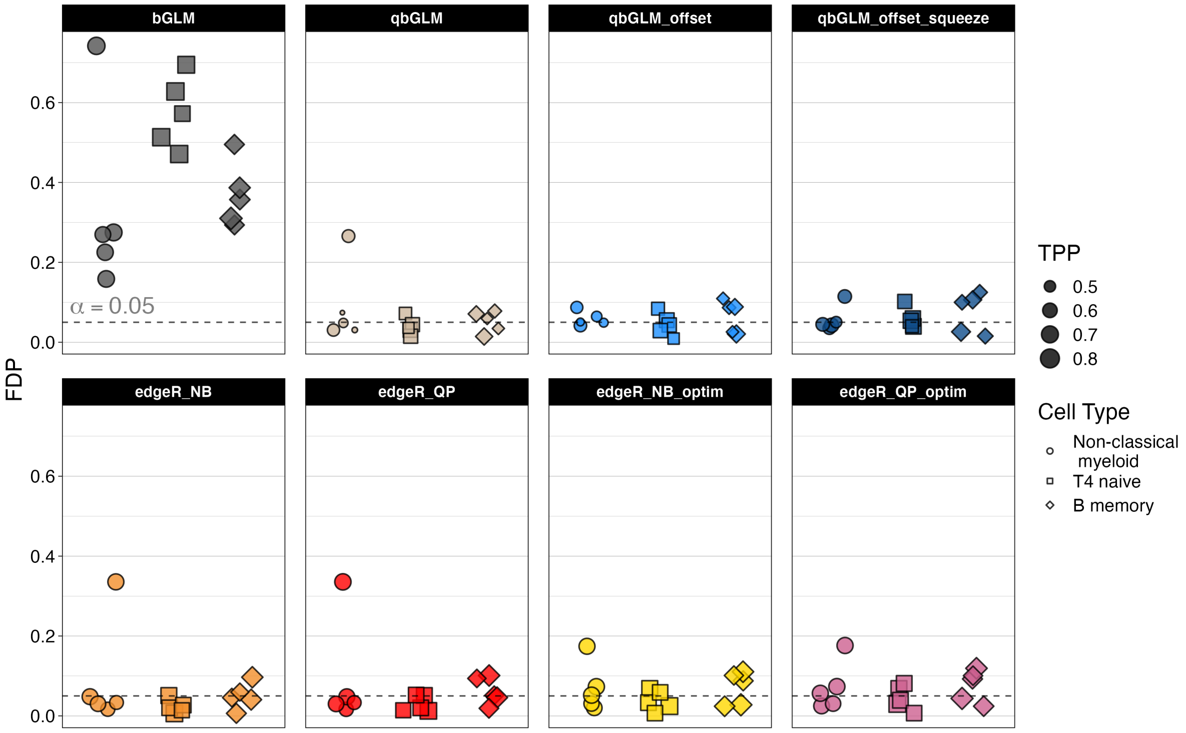


Supplementary figure 8: **Alternative visualization for the performance evaluation of eight differential detection methods on three simulated datasets.** The simulated data is based on the three different celltypes (non-classical myeloid cells, T4 naive cells and memory B cells) of the lupus dataset. The data was aggregated to the pseudobulk level, resulting in a test between two groups of five samples each. This plot is an alternative visualization for the results show in figure 2 of the main manuscript, which allows for showing the false discovery proportion (FDP) and true positive proportion (TPP) for the individual simulation replicates.

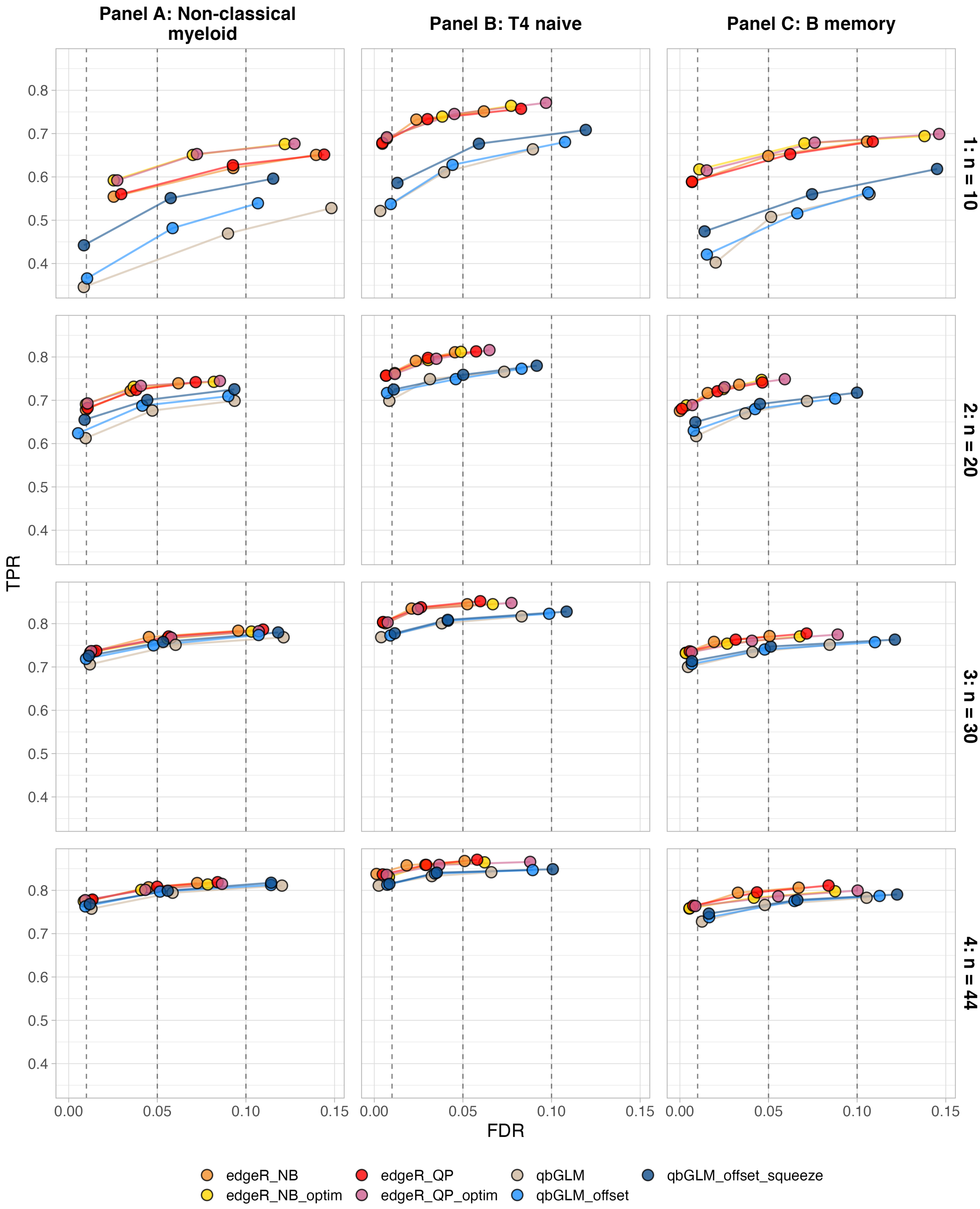


Supplementary figure 9: **Performance evaluation of eight differential detection methods on twelve simulated datasets.** The simulated data is based on the three different celltypes (non-classical myeloid cells, T4 naive cells and memory B cells) of the lupus dataset. The data was aggregated to the pseudobulk level, resulting in a test between two groups of five, ten, fifteen or twenty-two samples each (in the rows). Each curve visualizes the performance of each method by displaying the sensitivity of the method (true positive rate, TPR) with respect to the false discovery rate (FDR). The curves display averages over 5 replicates for each simulated dataset. The three circles on each curve represent working points when the FDR level is set at nominal levels of 1%, 5% and 10%, respectively.

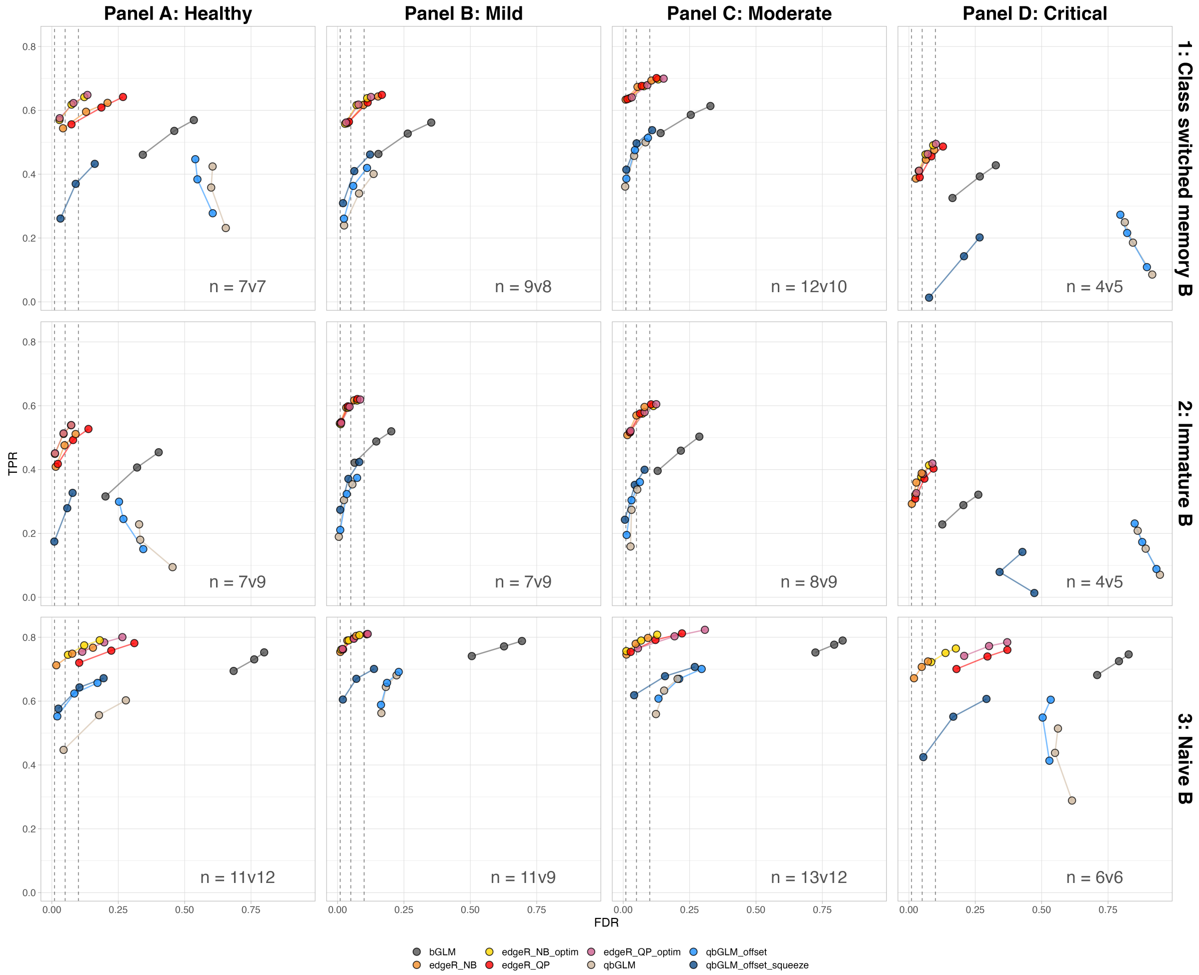


Supplementary figure 10: **Performance evaluation of eight differential detection methods on twelve simulated datasets.** The simulated data is based on the three different celltypes (class switched memory B cells, immature B cells and naive B cells, in the rows) of the covid dataset. The data was aggregated to the pseudobulk level for patients of the same disease severity (healthy, mildly disease, moderately diseased and criticall diseased patients in the columns). Each curve visualizes the performance of each method by displaying the sensitivity of the method (true positive rate, TPR) with respect to the false discovery rate (FDR). The curves display averages over 5 replicates for each simulated dataset. The three circles on each curve represent working points when the FDR level is set at nominal levels of 1%, 5% and 10%, respectively.

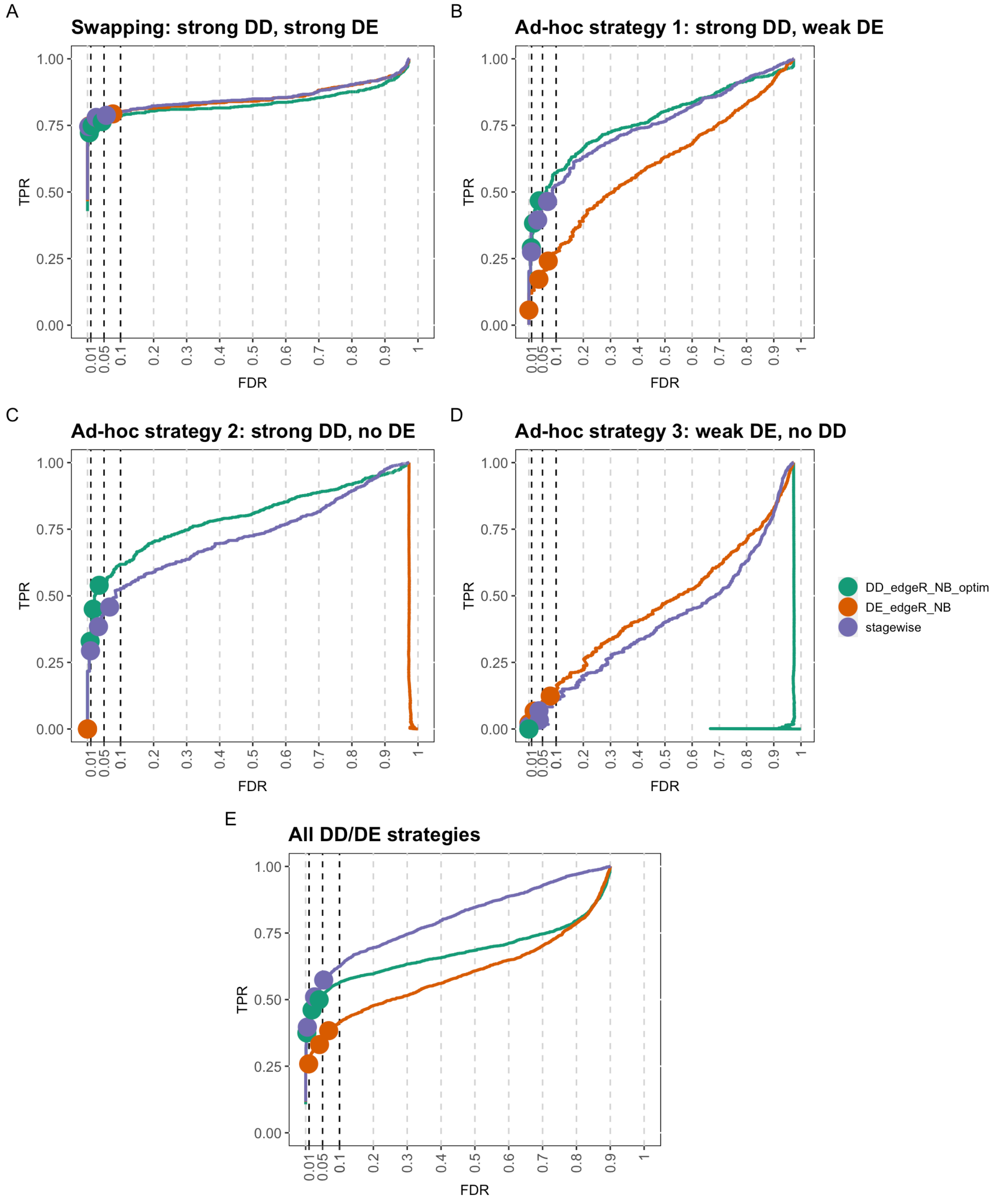


Supplementary figure 11: **Performance evaluation of differential detection, differential expression and stage-wise integrated workflows under different simulation strategies.** The starting point of these simulations is mock comparison on the non-classical myeloid cell from the lupus dataset, downsampled to a 5 versus 5 patients comparison. Next, we generated differential detection and/or differential expression signal according to four different simulation strategies. Performance curves for the differential detection, differential expression and stage-wise integrated workflows are displayed in green, orange and purple, respectively. **Panels A-D:** Performance evaluation on data with differential signal generated using the swapping strategy (panel A), the first ad-hoc simulation strategy (panel B) the second ad-hoc simulation strategy (panel C) and the third ad-hoc simulation strategy (Panel D). Note that the strategy construct panel A is the same as the strategy used to construct figure 2. **Panel E:** Performance evaluation in which signal generated using all four simulation frameworks is present at equal proportions. Here, the stage-wise analysis strongly outperforms the individual workflows, because for each gene with signal at least one of the two workflows is able to detect the differential signal.

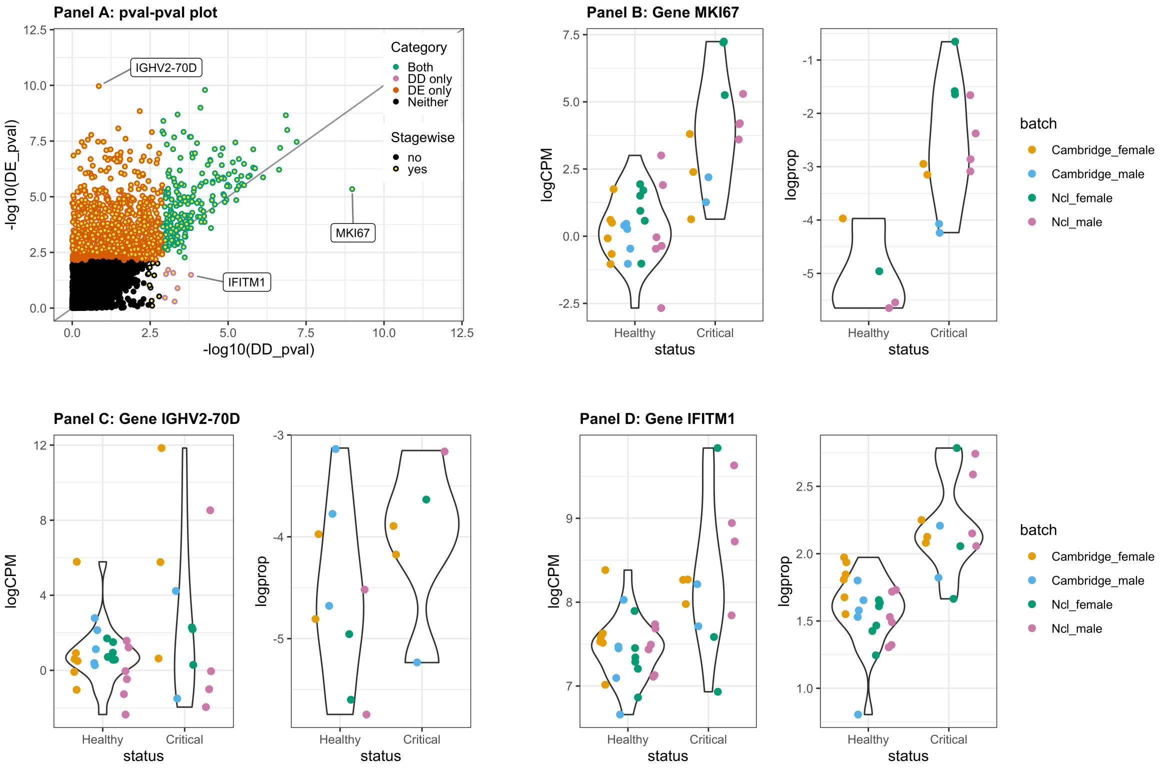


Supplementary figure 12: Complementarity of DE and DD analysis results for the comparison within naive B cells between healthy donors and critically ill COVID patients. Panel A: DE and DD analysis transformed p-values. The p-values from the DD analysis are displayed against the p-values of the DE analysis (upon -log10 transformation). Each dot represents the statistical significance of a single gene in the DE and DD analysis. Statistical significance was assessed at an FDR of 5%. Each is either both DE and DD (green), only DE (red), only DD (pink) or neither DE nor DD (black). Genes that passed the screening stage of the stage-wise testing procedure are indicated with a yellow center in the dot. Three genes are highlighted in this panel: MKI67, IGHV2-70D and IFITM1. These will be considered archetypes for genes that are both DE and DD, only DE or only DD, respectively. Panel B: Violin plots for gene MKI67. The left panel displays a measure of gene expression, while the right panel displays a measure for gene detection. In line with panel A, this gene has a clear signal of both DE and DD. Panel C: Violin plots for gene IGHV2-70D. In line with panel A, this gene has a strong DE signal, but no evidence for DD. Panel D: Violin plots for gene IFITM1. In line with panel A, this gene has a strong DD signal. Visually, a small shift in the average expression can also be observed, i.e. when visually comparing the average logCPM expression between healthy donors and COVID patients within each batch. This signal was not picked up by the DE analysis, likely because the large variability in the COVID patients and the relatively small sample size.

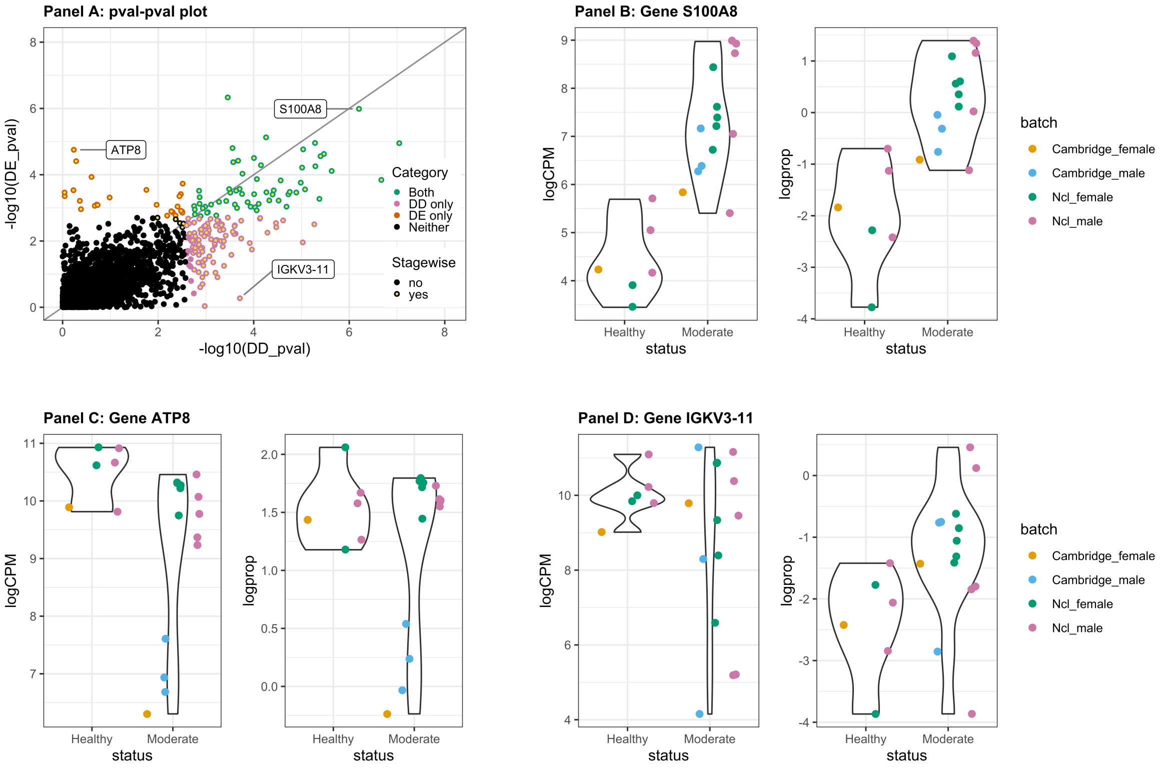


Supplementary figure 13: Complementarity of DE and DD analysis results for the comparison within unswitched memory B cells between healthy donors and moderately ill COVID patients. Panel A: DE and DD analysis transformed p-values. The p-values from the DD analysis are displayed against the p-values of the DE analysis (upon -log10 transformation). Each dot represents the statistical significance of a single gene in the DE and DD analysis. Statistical significance was assessed at an FDR of 5%. Each is either both DE and DD (green), only DE (red), only DD (pink) or neither DE nor DD (black). Genes that passed the screening stage of the stage-wise testing procedure are indicated with a yellow center in the dot. Three genes are highlighted in this panel: S100A8, ATP8 and IGKV3-11. These will be considered archetypes for genes that are both DE and DD, only DE or only DD, respectively. Panel B: Violin plots for gene S100A8. The left panel displays a measure of gene expression, while the right panel displays a measure for gene detection. In line with panel A, this gene has a clear signal of both DE and DD. Panel C: Violin plots for gene ATP8. In line with panel A, this gene has a strong DE signal. Visually, some evidence for DD also seems to be present, i.e., comparing the logCPM expression between healthy donors and COVID patients within each batch. This signal wass not picked up by the DE analysis, likely due to the small sample size. Panel D: Violin plots for gene IGKV3-11. In line with panel A, this gene has a strong DD signal. Visually, a small shift in the average expression can also be observed, i.e. when visually comparing the average logCPM expression between healthy donors and COVID patients within each batch. This signal was not picked up by the DE analysis, likely because the large variability in the COVID patients and the relatively small sample size.

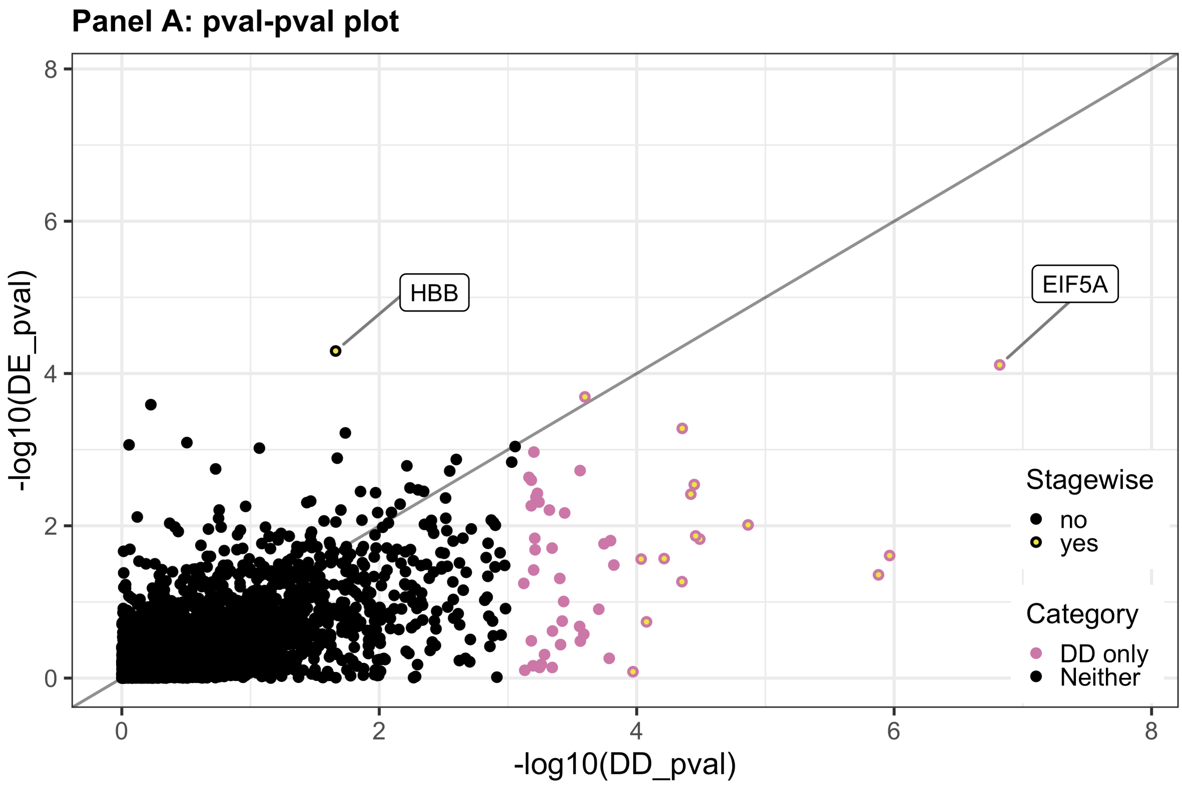


Supplementary figure 14: Complementarity of DE and DD analysis results for the comparison within unswitched memory B cells between healthy donors and critically ill COVID patients. Panel A: DE and DD analysis transformed p-values. The p-values from the DD analysis are displayed against the p-values of the DE analysis (upon -log10 transformation). Each dot represents the statistical significance of a single gene in the DE and DD analysis. Statistical significance was assessed at an FDR of 5%. Each is either both DE and DD (green), only DE (red), only DD (pink) or neither DE nor DD (black). Genes that passed the screening stage of the stage-wise testing procedure are indicated with a yellow center in the dot. Two genes are highlighted in this panel: EIF5A and HBB. These will be considered archetypes for genes that are both DE and DD, only DE or only DD, respectively.

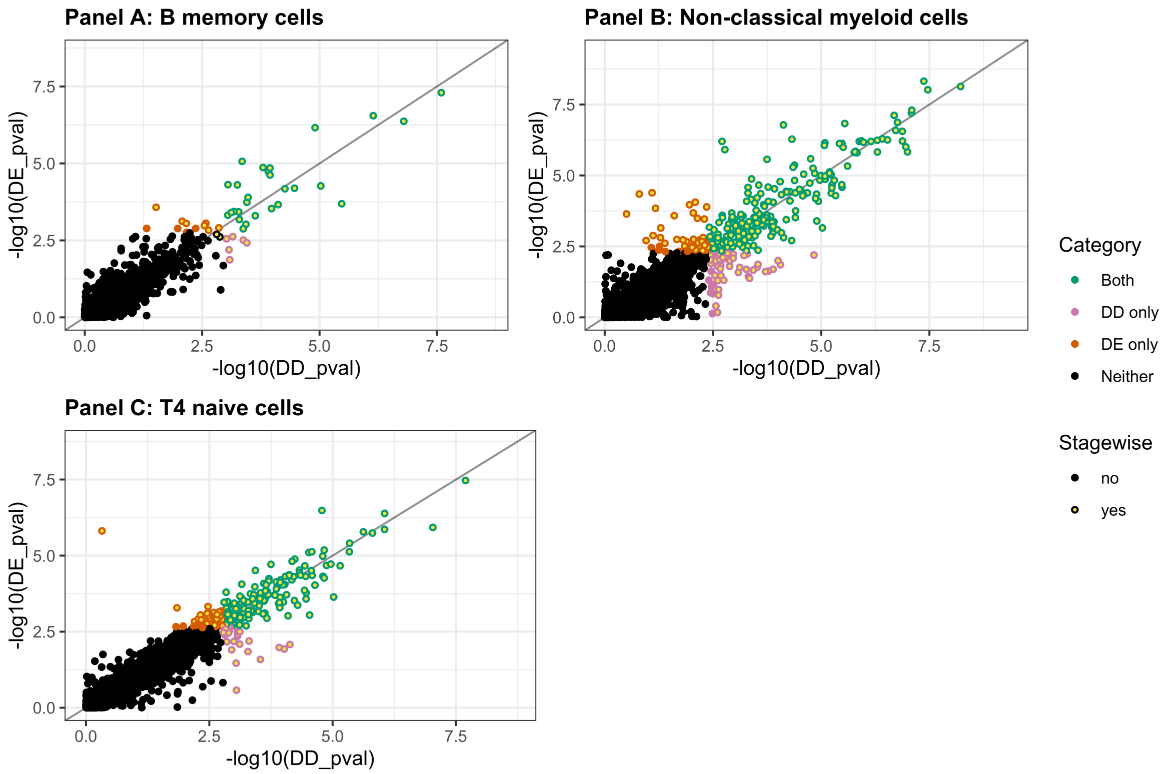


Supplementary figure 15: Complementarity of DE and DD analysis results for the lupus case study. Each panels displays the p-values from the DD analysis plotted against the p-values of the DE analysis (upon -log10 transformation). Each dot represents the statistical significance of a single gene in the DE and DD analysis. Statistical significance was assessed at an FDR of 5%. Each is either both DE and DD (green), only DE (red), only DD (pink) or neither DE nor DD (black). Genes that passed the screening stage of the stage-wise testing procedure are indicated with a yellow center in the dot. Panel A: B memory cell type. Panel B: Non-classical myeloid cell type. Panel B: T4 naive cell type.
